# Bridging genomes and peptidomes: hybrid sequencing reveals conserved bioactive peptides in crustaceans

**DOI:** 10.64898/2026.05.04.721987

**Authors:** Lauren Fields, Jiangrong Qin, Angel E. Ibarra, Kendra G. Selby, Tong Gao, Tina C. Dang, Haiyan Lu, Lingjun Li

## Abstract

Endogenous peptides are critical regulators of signaling and immunity but remain difficult to characterize in organisms with incomplete genomic annotation. We developed a hybrid discovery platform that integrates transformer-based *de novo* sequencing (Casanovo), neuropeptide-focused database searching (EndoGenius), and empirical false discovery rate estimation via NovoBoard. This pipeline enables confident identification of endogenous peptides while expanding coverage beyond conventional database-only or *de novo*-only approaches. Applied to neuroendocrine tissues from *Callinectes sapidus* and *Cancer borealis*, the workflow revealed numerous high-abundance novel peptides and provided structural and genomic support for their biological relevance. Notably, we report the first histone-2A–derived antimicrobial peptide in the *C. sapidus* and characterize naturally occurring sequence variants. We also identified unexpected peptide homologies between crustaceans and *Rattus norvegicus*, enabling annotation of conserved housekeeping proteins in sparsely annotated genomes. This hybrid platform establishes a scalable, open-source strategy for advancing neuropeptidomics and endogenous peptide discovery in emerging model organisms.

## Introduction

Endogenous peptides, such as antimicrobial peptides (AMPs) and neuropeptides, are key biomolecules that can be cleaved from protein precursors and exert unique biological activities to modulate physiological responses.^1,2^ The diverse and potent functionality of these peptides has made them attractive targets for biomarker and therapeutic development.^2^ Crustacean model systems have driven foundational discoveries in neuropeptide signaling and innate immunity, owing in part to their simple nervous system and lack of an adaptive immune system, which together make them ideal for discovery and evaluation of endogenous, bioactive peptides.^3–5^ For example, blue crabs (*Callinectes sapidus*), like the ones in this study, have long been revered for their production of AMPs, perhaps most famously the peptide Callinectin.^6^ Additionally, Jonah crabs (*Cancer borealis*) house the most well-characterized model system of pyloric rhythms and gastric mill neural circuits, fundamental to feeding behavior.^7^

Analysis of endogenous peptides presents several key challenges, particularly their low abundance *in vivo*, susceptibility to degradation, and poorly characterized cleavage patterns.^2^ Mass spectrometry (MS) is considered the gold standard for peptidomic analyses due to its sensitivity, throughput, and ability to derive sequence information from fragmentation spectra. In a typical peptidomic workflow, MS acquisition is followed by database searching, in which experimental spectra are matched to in silico fragmentation predictions derived from a supplied protein or peptide database. Searching input databases in a digest-free manner has been extensively applied for endogenous peptide profiling;^8^ however, cleavage prediction in this manner limits the ability to account for degraded forms of longer, mature peptides that have not been observed previously. Moreover, these search strategies present a unique complication for crustacean model systems due to many crustacean genomes only recently being assembled,^9–12^ thus making it difficult to provide a fully comprehensive endogenous peptide database for identification. Despite the tremendous advantages of using crustacean model systems for primary research of endogenous peptides, these analytical challenges have limited the ability to comprehensively characterize the crustacean peptidome.

Because crustacean peptidomic research remains far removed from genomic- or transcriptomic-level sequence characterization, scientists have relied on highly untargeted methods, such as *de novo* sequencing, to identify novel, potentially bioactive endogenous peptides.^7,13–18^ *De novo* sequencing is a well-established approach for identification of peptides from MS data, but historically proved cumbersome due to the extensive computational time requirements. With the advent of higher resolution MS instrumentation and faster computational capabilities, *de novo* sequencing has experienced a resurgence with several recently-released platforms such as DiNovo,^19^ PepNet ^20^, GraphNovo ^21^, and Casanovo.^22,23^ Generally, these tools are trained on model datasets of target and decoy spectra to estimate peptide sequences by recognizing the correlation between patterns of fragment ions and amino acid sequences. *De novo* sequencing offers the ability to make peptide identifications without a prior database, making it useful when databases or genomic annotation is limited, as is the case for many crustacean species. Notably, *de novo* sequencing is not intended to completely replace database searching, but rather to complement it; however, principled frameworks for integrating the two approaches remain limited, particularly for endogenous peptides.

In this work, we define a hybrid platform composed of Casanovo for *de novo* sequencing,^24,25^ and our novel database searching platform, EndoGenius, tailored specifically for endogenous peptides.^26^ We hypothesized that this two-step strategy of *de novo* sequencing and database searching could achieve a more comprehensive representation of the crustacean peptidome by offering detection of novel endogenous peptides not present in current sequence databases. This theory was rooted in proteomics software that models this approach; ^27^ however, their applicability to non-digested peptides is limited, thus prompting our investigation. Data were acquired from control (*i.e.*, acclimated and unperturbed) *C. sapidus* neuroendocrine tissues as a validation dataset. Mass spectra were processed via a standard database search in EndoGenius, and in parallel, subjected to a rigorous *de novo* sequencing pipeline composed of five key steps: (1) generation of decoy spectra using NovoBoard, (2) *de novo* sequencing via Casanovo, (3) reentry of the predictions into NovoBoard for filtering to 5% false discovery rate (FDR), (4) searching the confident predictions using EndoGenius, and finally (5) integration of the *de novo* results with those identified via database searching (**Fig. 1**). This hybrid method was then applied to *C. borealis* neuroendocrine tissue from fed crabs. Through this method, we discovered the first reported histone-2A-derived AMP in *C. sapidus*, defining its organization within the genome and characterizing a naturally occurring sequence variant. Additionally, we discovered novel, high-abundance crustacean peptides which showed homology with *Rattus norvegicus* (Sprague-Dawley rat), leveraging the more complete annotation of the rat proteome to define key housekeeping proteins in *C. borealis* for the first time. Our results establish a hybrid pipeline for discovery and characterization of endogenous peptides in organisms with limited genomic resources.

**Figure 1:**
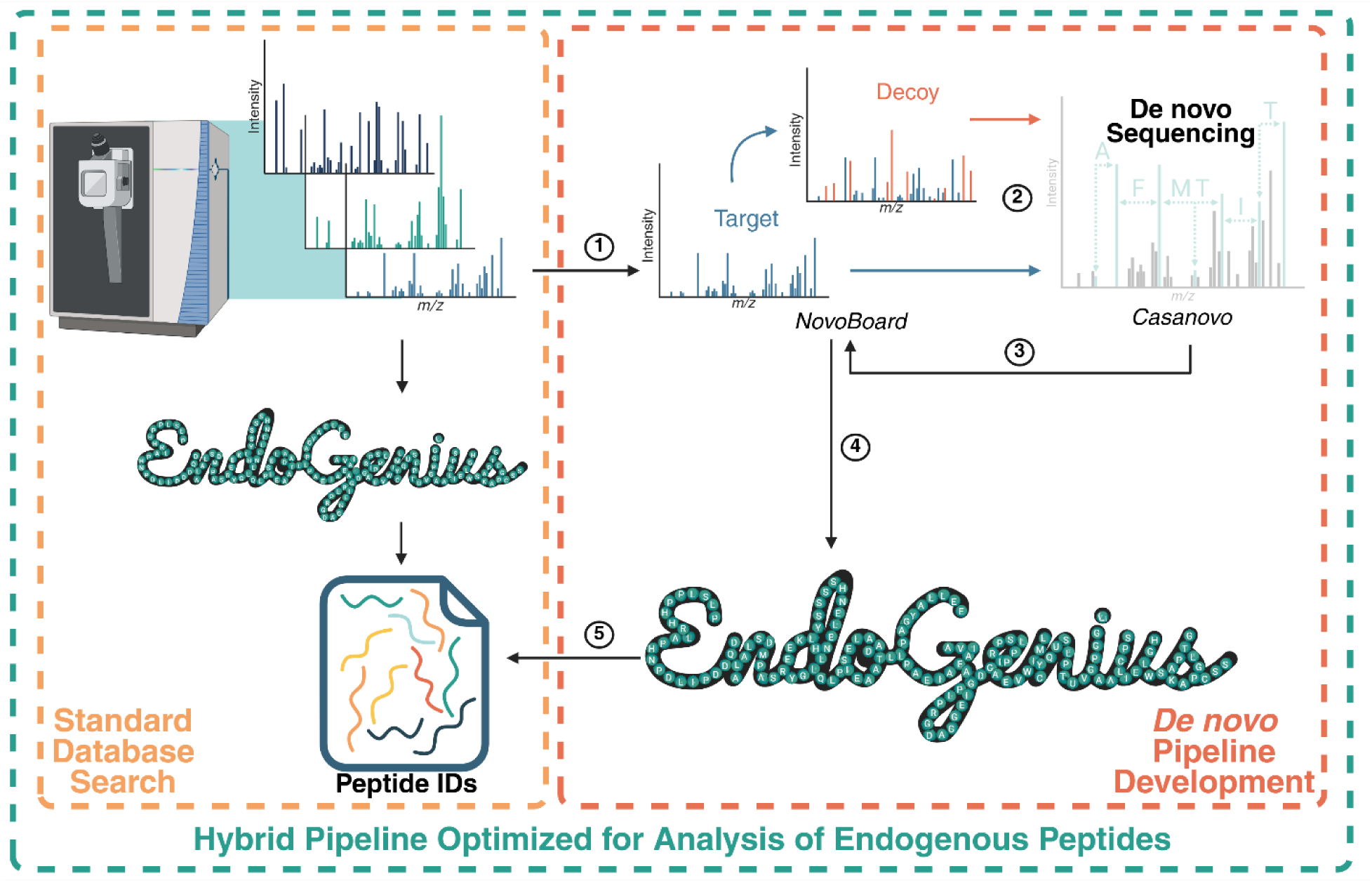
S**c**hematic **of hybrid analytical pipeline for characterization of endogenous peptides.** Following MS acquisition, spectra are processed directly in EndoGenius to yield identifications according to a reference database (left, orange box). Simultaneously, data are also processed through the *de novo* component of the pipeline (right, red box), which is composed of five general steps: (1) spectra are supplied to NovoBoard for generation of decoy spectra, (2) target and decoy spectra are transferred to Casanovo for *de novo* sequencing, (3) predicted sequences are returned to NovoBoard for filtering at 5% false discovery rate (FDR), (4) confident *de novo* results are searched using EndoGenius, and (5) the final results are integrated with the standard database search. Integration of both components yields the final hybrid pipeline for endogenous peptidomics (teal box).

## Results

### Establishment and justification of hybrid analytical pipeline for endogenous peptides

Hybrid data analysis pipelines integrate *de novo* sequencing with database searching, enabling both confident identification and novel discovery of peptides from MS data. This approach is particularly advantageous for endogenous peptides, which exhibit high sequence similarity and noncanonical cleavage patterns,^8^ complicating isoforms discrimination. Given that key crustacean model organisms such as *C. sapidus* and *C. borealis* only recently had their genomes assembled and remain sparsely annotated,^9,28^ a hybrid pipeline is well suited to expand peptidomic coverage in these systems.

We have previously shown that even refined hybrid pipelines require optimization and post-analysis investigation to account for unique qualities of endogenous peptides, including high sequence homology and a wide length distribution.^29^ It is therefore important to construct a hybrid analytical pipeline specialized for analysis of endogenous peptides like AMPs and neuropeptides that requires less manual optimization than existing approaches. Importantly, all components of this pipeline are open source, ensuring availability to all researchers. In this work, we have integrated our database searching platform, EndoGenius, which is optimized for endogenous peptide analyses,^8,26^ with Casanovo, an open-source platform for *de novo* sequence analysis (**Fig. 1**).^25,30^

Casanovo was selected for its accuracy, scalability, and methodological transparency. Briefly, it uses a transformer-based architecture to directly model the relationship between fragment ion patterns and amino-acid sequences, enabling faster inference and improved accuracy relative to earlier de novo approaches.^30^ Notably, Casanovo offers several pretrained models optimized for different acquisition settings, including tryptic and enzymatic pretreatments, enabling flexible model selection without retraining.^24,25^ We posited that the large-scale mixed-enzyme training could be generalized to endogenous peptides that often deviate from canonical enzymatic digestion rules.

Like most *de novo* sequencing tools, Casanovo does not provide intrinsic false discovery rate control because there is no database from which to derive decoy sequences. To address this, we employed NovoBoard, which generates paired decoy spectra directly from each experimental MS² spectrum using controlled peak deletion and noise injection.^31^ This design preserves the global statistical structure of the data while removing peptide-specific fragmentation information, producing decoys that are biophysically plausible but do not correspond to real peptide sequences. Because NovoBoard operates independently of sequence databases, it ensures that validation reflects instrument-level and chemistry-specific characteristics of the spectra rather than assumptions about the underlying proteome. In addition to generating the decoy spectra, NovoBoard enables empirical estimation of FDR in *de novo* predictions.^31^ Previous benchmarking has demonstrated that a 50% peak-removal setting works best for Casanovo,^31^ and was therefore used for workflow validation and feeding applications in this study.

The proposed pipeline was first validated using control (*i.e.*, acclimated and unperturbed) blue crab neuroendocrine tissues, including the brain and thoracic ganglion (TG) as well as the paired pericardial organs (PO), sinus glands (SG), and commissural ganglia (CoG). MS data were subjected to a standard database search against our in-house developed crustacean peptide database^29^ using EndoGenius (**Fig. 1, left**). In parallel, paired decoy spectra for each target spectrum were generated using NovoBoard, followed by *de novo* sequencing via Casanovo (**Fig. 1, right, steps 1 & 2**). *De novo* results were filtered to the equivalent of 5% FDR using the NovoBoard FDR framework (**Fig. 1, right, step 3**). Identifications passing this threshold were then subjected to a second iteration of database searching, filtering to a confident EndoGenius score of 1000 (**Fig. 1, right, step 4**).^26^ Thus, the final results comprise both known, database-identified peptides, and novel, putative endogenous peptides (**Fig. 1, right, step 5**). The same workflow (**Fig. 1**) was then applied to the corresponding tissues from fed Jonah crabs to extend these insights to feeding, a physiological process with well-established neuropeptidomic regulation.

### Assessment of de novo predictions

Following conceptualization of the hybrid pipeline (**Fig. 1**), its performance was assessed using a control blue crab dataset as described above. First, the quality of the decoy spectral outputs from NovoBoard was assessed to confirm the reliability of initial *de novo* predictions and downstream FDR estimates. It is anticipated that the decoy spectra generated by NovoBoard should yield score distributions matched to those of the target inputs. Considering the spectral characteristics of the target data, most fragment ions are low mass (**Fig. 2a, Supplementary Fig. 1**), with their intensities being approximately normally distributed, with a slight tail toward higher intensities (**Fig. 2b, Supplementary Fig. 2**). Kernel density estimates showed that the decoy spectra were highly similar to those of the target spectra with regards to not only fragment intensity (**Fig. 2b, Supplementary Fig. 1**) and *m/z* distribution (**Fig. 2b, Supplementary Fig. 2**), individually, but also in the distribution of fragment intensities per *m/z* (**Fig. 2c, Supplementary Fig. 3**). Together, these data confirm that NovoBoard produces chemically plausible decoy spectra that faithfully capture the statistical properties of the experimental data, enabling robust calibration of Casanovo confidence scores and rigorous per-file FDR control.

**Figure 2.**
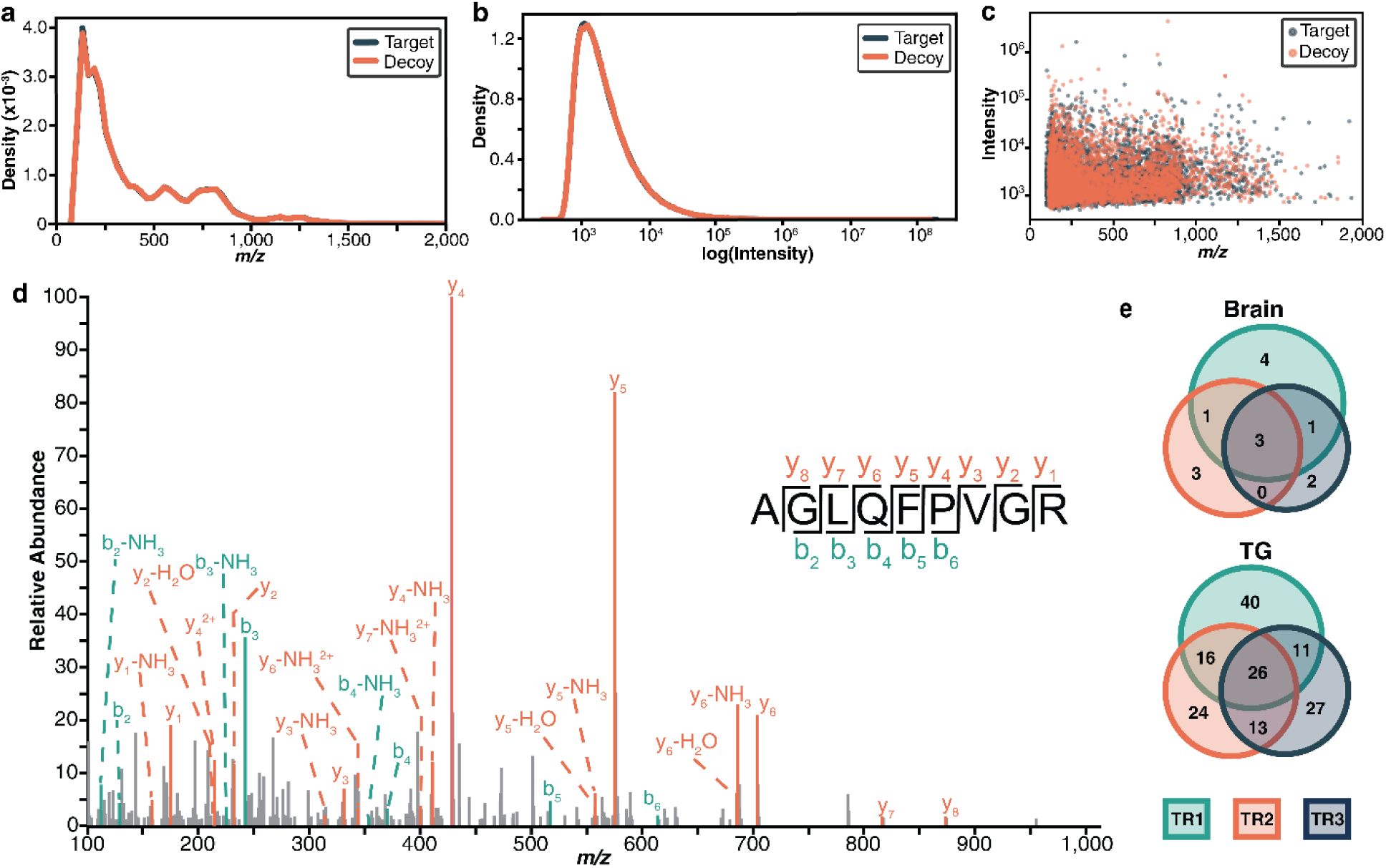
:Confidence assessment of hybrid pipeline for peptidomic analyses. Comparison of target (dark teal) spectral characteristics and NovoBoard-generated decoy (coral) spectra derived from them, including a) fragment *m/z*, b) fragment intensity, and c) intensity vs *m/z*. d) Fragmentation pattern of a representative neuropeptide identified via *de novo* sequencing, and subsequently validated in EndoGenius. e) Overlap of identifications from *de novo* sequencing of three technical replicates (TR) in the brain and thoracic ganglion (TG).

Following validation of decoy spectral quality, spectral coverage of the predicted peptides was evaluated to further validate the de novo results. Manual inspection of mass spectra for representative peptides such as AGLQFPVGR revealed extensive sequence coverage, demonstrating strong experimental support for the predicted sequences (**Fig. 2d**). This provided evidence that Casanovo can generalize to non-tryptic experimental conditions despite not having been trained on such data. Together with the decoy validation described above, these findings support the use of Casanovo and NovoBoard for neuropeptidomic applications.

Finally, the *de novo* prediction results from the three technical replicates for each tissue were assessed for reproducibility of prediction of novel, putative endogenous peptides. This step is important, as high identification confidence does not necessarily indicate biological relevance. Identifications from analysis of cardiac-related tissues including the TG (**Fig. 2e**) and PO (**Supplementary Fig. 4**) showed substantial overlap across triplicate analyses. Conversely, the brain (**Fig. 2e**), SG, and CoG (**Supplementary Fig. 4**) demonstrated minimal overlap, with no shared peptides for the SG triplicates. This pattern is consistent with the current state of crustacean peptidomics: neural tissues such as the brain and STNS (including the CoG) have been extensively characterized, leaving fewer unidentified peptides for de novo sequencing to capture. Novel identifications in these tissues are therefore rarer and more stochastically sampled across replicates. By contrast, cardiac-associated tissues such as the TG and PO remain comparatively underexplored, providing a larger pool of uncharacterized peptides that are more consistently detected across replicates. Though there is notable variability in the number of *de novo* predictions from each replicate (**Figs. 2e, Supplementary Fig. 4**), this is likely the result of performing data-dependent acquisition, which frequently overlooks low-abundance analytes.^2^ These results highlight the present gap in complete neuropeptidomic characterization for neuroendocrine-cardiac tissues, as well as the importance of replicate-level assessment for evaluating novel peptide predictions.

### De novo sequencing expands support for known peptides

Traditional database searches are not fully compatible with peptidomic analyses due to non-canonical cleavage patterns that can cause peptides to be overlooked. Hybridizing database searching with de novo sequencing can help recover these missed identifications. Moreover, it is critically important to use a database search tool specifically designed for endogenous peptides, capable of capturing their nuances, including the notable prevalence of b-ions in non-digested peptide spectra, including the notable prevalence of b-ions for non-digested peptides.^26^ Following a search of the *de novo* results from the blue crab validation dataset by EndoGenius, the proportion of b- and y-ion coverage was compared to that of the direct database search in EndoGenius (**Fig. 3a**). Integration of *de novo* predictions with EndoGenius database search results revealed a more balanced distribution of b- and y-ion coverage (**Fig. 3a**). The shift toward greater y-ion coverage is likely the result of Casanovo having been trained on enzymatically digested datasets,^24^ of which tryptic peptides are known to produce mostly y-ions.^32^ Despite detection of more y-ions, this pipeline still retains a slight dominance of b-ions (**Fig. 3a, Supplementary Fig. 5**), as we have observed previously for digest-free experiments. Though the relationship between b-ion prevalence and endogenous peptides is not yet fully understood, it appears to be a characteristic feature of digest-free analyses. That *de novo* peptides generally maintaining this pattern further underscores the flexibility of Casanovo and supports its use in this hybrid pipeline, as this is the first application of Casanovo for *de novo* sequencing of non-digested peptides.

**Figure 3:**
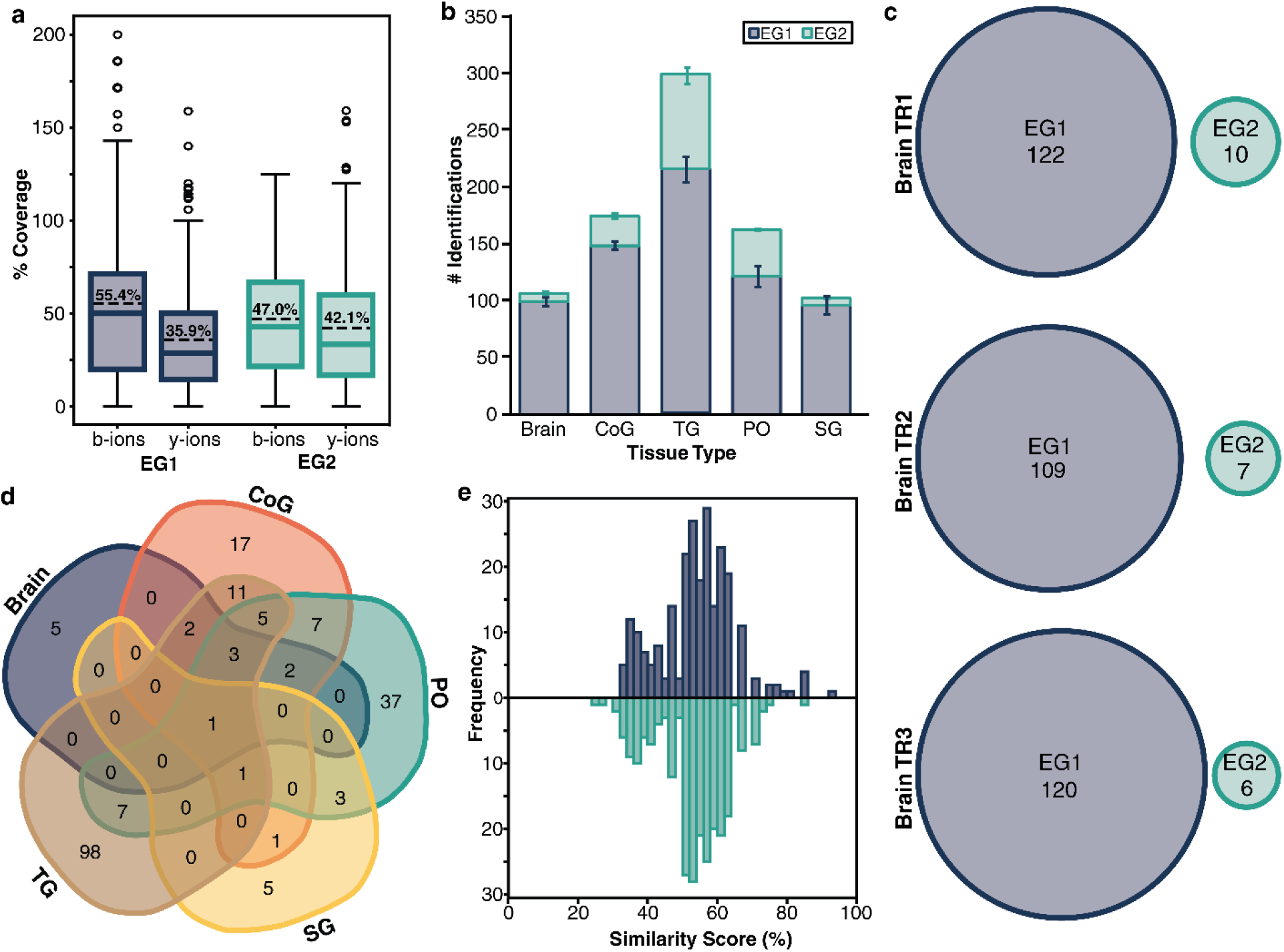
E**v**aluation **of putative endogenous peptide identifications from hybrid analytical pipeline. a)** Percent sequence coverage by b- and y-ions for direct database searching of spectra via EndoGenius (EG1, dark blue) and hybrid results containing database-identified and *de novo* peptides validated in EndoGenius (EG2, teal). **b)** Number of neuropeptide identifications afforded by database (EG1, dark blue) and hybrid (EG2, teal) results. Error bars represent standard deviation (*n* = 3) in identification counts across technical replicates. **c)** Overlap of brain identifications in EG1 (dark blue) and EG2 (teal) for three technical replicates (TR), revealing zero overlap. **d)** Overlap of validated *de novo* sequencing identifications (EG2) across five tissues: thoracic ganglion (TG), commissural ganglia (CoG), pericardial organs (PO), brain, and sinus glands (SG). **e)** Cumulative multiple sequence alignments between final results from the EG2-level of the hybrid pipeline and an in-house crustacean neuropeptide database^26^ (top, dark blue) and database of putative neuropeptides recently proposed^2^ (bottom, teal).

Having validated the *de novo* predictions, we next examined the identifications themselves. Direct database searching of the spectra by EndoGenius revealed approximately 100-200 identifications arising from each tissue, with most identifications from the TG, the largest tissue (**Fig. 3b, EG1**). These counts were expanded through incorporation of results from an EndoGenius search of the *de novo* results, with an increase by as many as 128 peptides in the TG (**Fig. 3b, EG2**). The brain and SG showed only moderate increases (**Fig. 3b, EG2**), agreeing with the raw *de novo* results (**Supplementary Fig. 4**). Both searches (*i.e.*, with and without *de novo* result incorporation) applied to triplicate injections revealed similar numbers of identifications for each search across tissue types (**Fig. 3c, Supplementary Fig. 6**). Moreover, comparison of the output of these two searches in the brain, for example, reveals no overlap (**Fig. 3c**) as would be expected. Several tissues demonstrated minor (five or less peptides) overlap (**Supplementary Fig. 6**), representing modified peptides which were not filtered from the *de novo* results. These results highlight the tissue-independent manner by which *de novo* sequencing expanded the number of peptide identifications through introduction of peptides that were unidentifiable by EndoGenius alone. Extending the tissue-based comparison of these results revealed that most of the novel peptides found in the PO and TG were found uniquely in these tissues, with approximately 58% and 77%, respectively, arising from only one of these tissues (**Fig. 3d**). The notable presence of novel, tissue-specific putative peptides further underscores the current gap in crustacean peptidomics surrounding investigation of neuroendocrine tissues with roles in the cardiac system (**Figs. 2b, 3b**).

The *de novo* results were also aligned with established neuropeptide databases to assess their biological plausibility as putative bioactive peptides. After removing known crustacean neuropeptides from the de novo results, the remaining sequences were compared to our in-house developed crustacean database (**Fig. 3e, top**) and a larger general neuropeptide database (**Fig. 3e, bottom**) in a tissue-independent manner. Comparison with either database revealed similar distributions, with most peptides showing 50-60% sequence similarity with entries in the respective database. Thus, Casanovo *de novo* sequencing enabled the identification of not only crustacean neuropeptides, but also neuropeptides originating from other model organisms, suggesting that many of these novel peptides share evolutionary homology with neuropeptides from other organisms, highlighting sequence conservation across the animal kingdom.

From these validation results, one novel crustacean peptide appeared in all five tissues: AGLQFPVGR (**Fig. 3d**). Investigation of the relevance of this peptide revealed this peptide corresponds to a fragment of Histone H2A type 1-B/E, which has been extensively characterized in mammals.^33^ This peptide was recently identified as an antimicrobial peptide with antioxidant activity in the black soldier fly (*Hermetia illucens L.*).^34^ Another study reported AGLQFPVGR from insects as having anti-obesity activity and protecting against inflammation, and is currently being studied as a potential therapeutic for metabolic dysfunction-associated steatotic liver disease.^35^ Of particular relevance, this peptide represents a portion of the 38-mer AMP Sphistin (MAGGKAGKDSGKAKAKAVSRSAR<u>AGLQFPVGR</u>IHRHLK, previously characterized in mud crabs (*Scylla paramamosain*). Sphistin has been reported as having Gram-positive and Gram-negative activity, as well as activity against yeast, with a mechanism of action that entails permeation of cell walls.^36^ Histone-derived antimicrobial peptides (HDAPs) such as Sphistin have only recently been discovered in crustaceans, with the current catalog of these peptides limited to just identifications from a few crabs, most recently the blue swimmer crab, *Portunus pelagicus*.^37^ To our knowledge, this is the first report of this HDAP in the blue crab, *Callinectes sapidus*.

The complete 64-mer HDAP sequence in C. sapidus was obtained by aligning the identified peptide against the *C. sapidus* genome, inclusive of the AGLQFPV hallmark motif.^9^ We observed the high frequency with which the HDAP is encoded within *C. sapidus*, with the complete 64-mer sequence encoded on chromosomes 2, 4, 5, 6, 7, 22, and 50 (**Fig. 4a**). Only one sequence was identified as an exact match on chromosome 7, while the remaining six chromosomes all encoded an identical variant containing four residue substitutions: V14I, L39M, T43A, and I46V (**Fig. 4a**). Notably, two of the four replacements, V14I and I46V, are isomeric residues which cannot be differentiated by MS^2^. These results underscore the high degree of conservation of this HDAP, which has likewise been reported elsewhere.^38^

**Figure 4:**
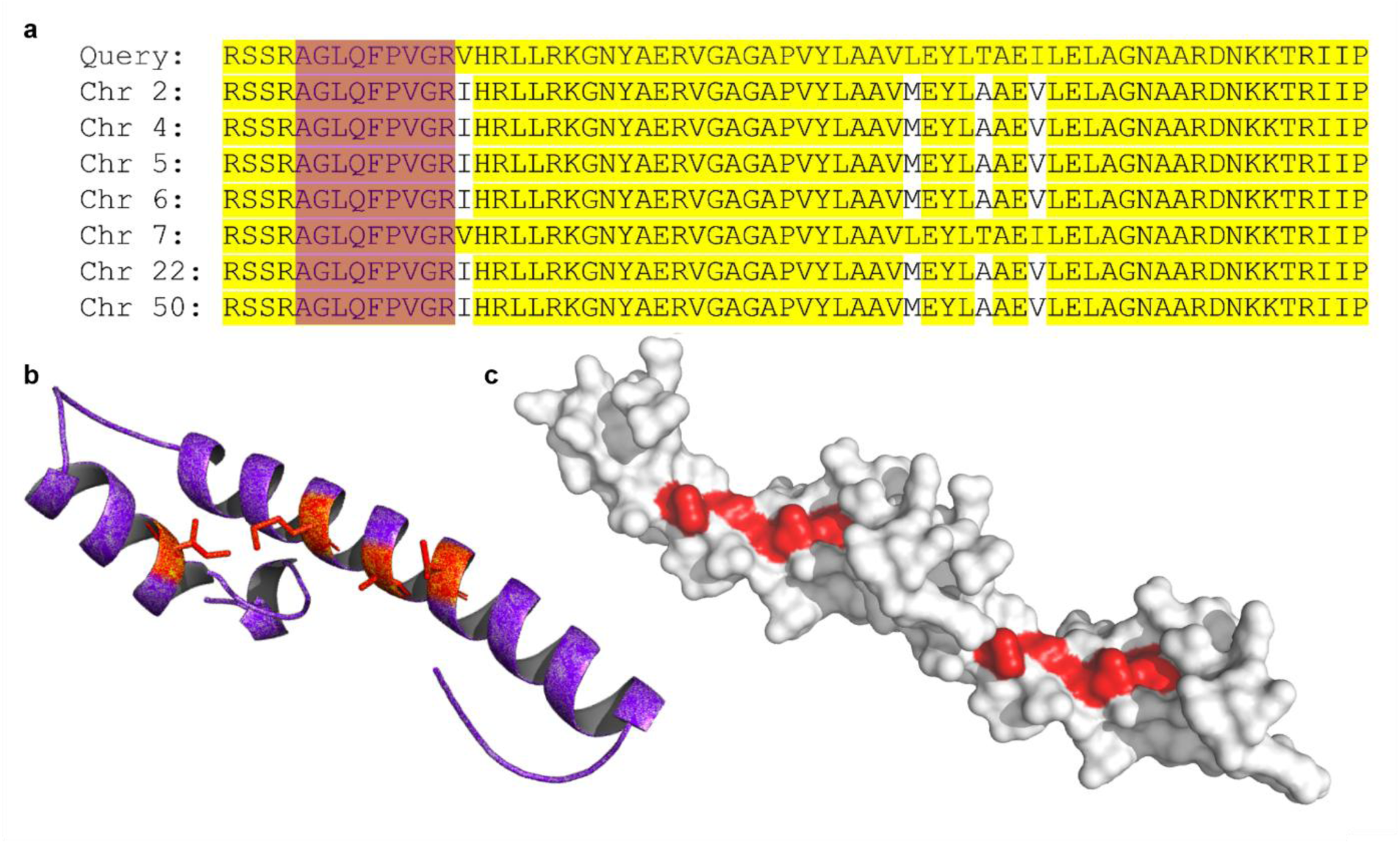
G**e**nomic **organization and predicted structure of histone-2A-derived peptides in *C. sapidus*. a)** Chromosomal sequence alignment for histone-derived antimicrobial peptide (HDAP), with conserved regions highlighted in yellow and the peptide identified in all tissues highlighted in magenta. **b)** Superimposed predicted structures of the query (Chr 7) and variant (Chrs 2, 4, 5, 6, 22, and 50) peptide sequences, showing near-identical backbone conformations. Substituted residues in the variant form are highlighted in red. **c)** Surface mapping of peptide, with red denoting residue deviations which were also highlighted in red in **Fig. 4b**.

Considering the structural impacts of these residue replacements, we constructed a predicted structure of these HDAP sequences.^39–41^ The exact match found on chromosome 7 was superimposed with the sequence variant containing V14I, L39M, T43A, and I46V substitutions, which was the same for all other chromosomes (**Fig. 4a**). Both sequences adopt the same overall fold, characterized by prominent helical structure (**Fig. 4b**), a canonical feature of many AMPs.^42^ Notably, when evaluating the surface of the aligned peptide, all four variant residues co-localize on the peptide surface despite being non-contiguous in the primary sequence (**Fig. 4c**). These results underscore conservation of this HDAP, while also suggesting that these variants may exhibit altered bioactivity, and warranting further investigation of their antimicrobial or binding properties.

The abundance of novel peptides was compared to that of their database-identified counterparts. Novel identifications spanned a wide dynamic range, with peptides detected across low, mid, and high intensity ranges, most notably in the TG (**Fig. 5a**), PO (**Fig. 5b**), and CoG (**Fig. 5c**). The brain (**Fig. 5d**) contributed few novel peptides, consistent with its extensive prior characterization. The SG (**Fig. 5e**) similarly yielded few novel identifications, likely reflecting its small size and limited peptide diversity. The broad dynamic range of novel peptides across other tissues highlights the capacity of de novo sequencing to expand peptidomic coverage beyond what is achievable by database searching alone.

**Figure 5.**
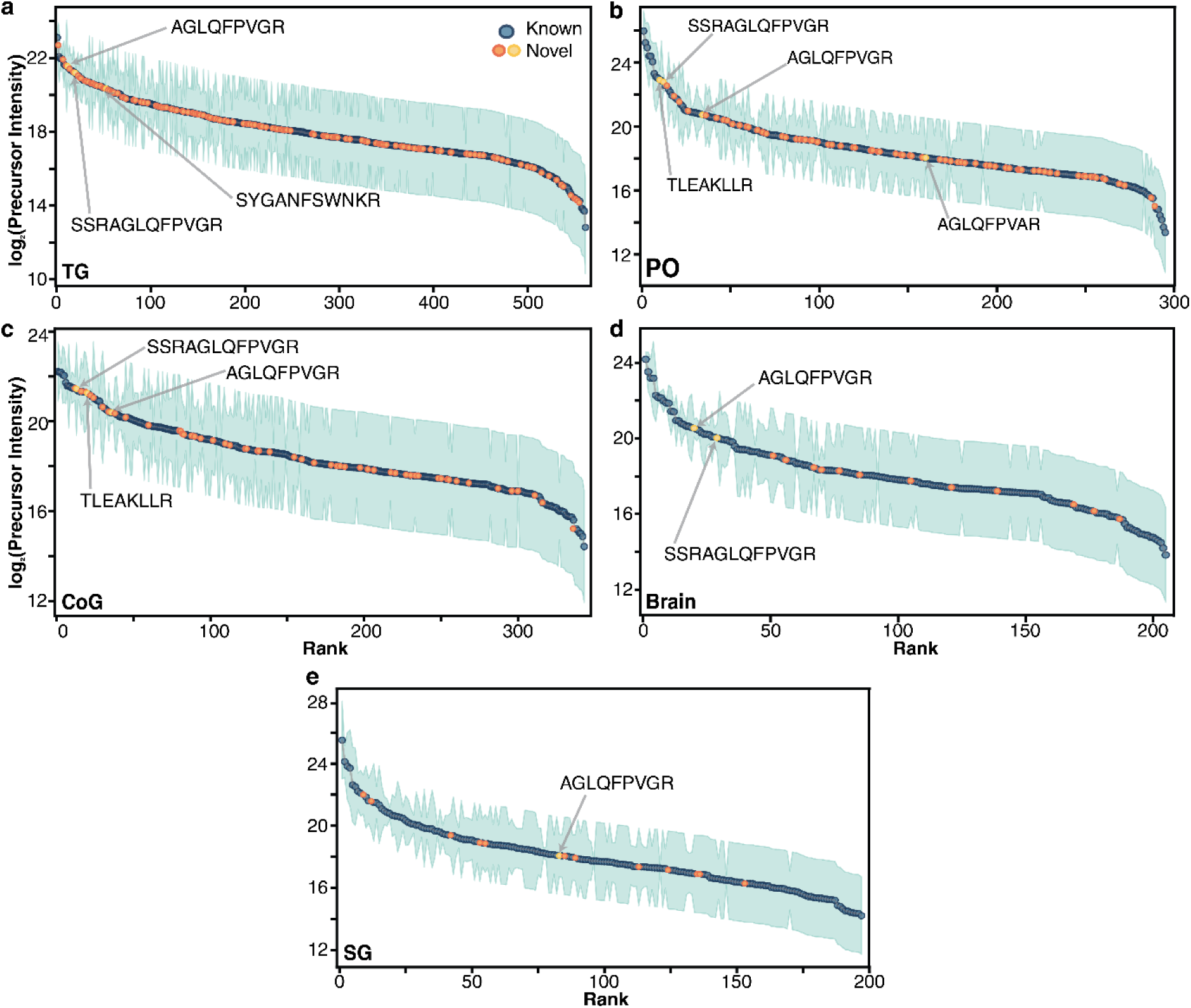
:Ranking of peptides identified in three technical replicates from crustacean neuroendocrine tissues. Novel (orange) and known (blue) peptides are shown for the a) thoracic ganglion (TG), b) pericardial organ (PO), c) commissural ganglia (CoG), d) brain, and e) sinus glands (SG). Novel peptides which were further investigated are highlighted in yellow.

Comparison of relative intensity across tissues revealed tissue-specific abundance patterns for novel peptides, such as the previously discussed HDAP peptide AGLQFPVGR, found in all five tissues. In the TG, this peptide ranked third among novel peptides and tenth overall (**Fig. 5a**). In other tissues, AGLQFPVGR ranked 34th, 36th, 20th, and 83rd overall in the PO (**Fig. 5b**), CoG (**Fig. 5c**), brain (**Fig. 5d**), and SG (**Fig. 5e**), respectively (**Table 1**), suggesting tissue-dependent variation in expression. By comparison, an extended form of this peptide, SSRAGLQFPVGR, was identified in all tissues except the SG, ranking 18th, 12th, 13th, and 29th overall in the TG (**Fig. 5a**), PO (**Fig. 5b**), CoG (**Fig. 5c**), and brain (**Fig. 5d**), respectively (**Table 1**). Though SSRAGLQFPVGR extends the previously discussed HDAP by three N-terminal residues, its intra- and inter-tissue relative abundance pattern is notably different, likely reflecting differential regulation of endogenous peptides and their isoforms across tissues.

**Table 1.** Abundance ranking and predicted bioactivity of top de novo-identified peptides.

| Peptide | Abundance Rank |  |  |  |  | AMP Potential<sup>a</sup> | Neuropeptide Potential<sup>b</sup> |  |
| --- | --- | --- | --- | --- | --- | --- | --- | --- |
|  | Brain | CoG | PO | SG | TG | DBAASP<sup>c</sup> | NeuroPred-FRL<sup>c</sup> | NeuroPP<sup>c</sup> |
| AGLQFPVGR | 20 | 36 | 34 | 83 | 10 | non-AMP | 0.881 | 0.85874 |
| SSRAGLQFPVGR | 29 | 13 | 12 |  | 18 | non-AMP | 0.614 | 0.9968 |
| STLTSRELQTAVR | 169 | 22 | 18 |  | 30 | non-AMP | 0.23 | 0.69954 |
| AKWDAWNSR |  | 12 | 48 |  | 2 | non-AMP | 0.83 | 0.96673 |
| TLEAKLLR |  | 20 | 10 |  |  | AMP | 0.67 | 0.56155 |
| FFVHWNR |  | 123 | 14 | 55 | 14 | AMP | 0.435 | 0.26045 |
| GSFMSVESLSR |  | 34 |  |  | 7 | non-AMP | 0.416 | 0.99678 |
| SNTTALAEAWAR |  |  |  |  | 20 | non-AMP | 0.099 | 0.65732 |

<sup>a</sup>Antimicrobial peptide (AMP) potential was evaluated via the Database of Antimicrobial Activity and Structure of Peptides (DBAASP).<sup>48</sup>
<sup>b</sup>Potential for neuropeptide activity was assessed by NeuroPred-FRL<sup>49</sup> and NeuroPP.<sup>50</sup>
<sup>c</sup> Green highlight indicates predicted AMP or neuropeptide activity by the respective tool.

Among other high-abundance peptides, several previously reported roles in diet and metabolism. The peptide AGLQFPVAR was identified in the PO and has been shown to have lipid-inhibitory capabilities as well as antimicrobial activity.^43^ This peptide differs from the previously discussed HDAP AGLQFPV<u>G</u>R by only one amino acid in the penultimate position

from the C-terminus, highlighting the diverse functionality of closely related peptide isoforms. Additionally, SYGANFSWNKR was found in the TG, which was previously identified in a study of cardiac protein changes following dietary soybean oil intake, with reported functions related to the electron-transport chain.^44^

While other high-abundance peptides lacked supporting evidence in the literature, we assessed all novel neuropeptides for activity as both neuropeptides and antimicrobial peptides. Antimicrobial peptides share many properties with neuropeptides,^8^ and indeed some peptides exhibit both antimicrobial and neuromodulatory activity (*e.g.*, pituitary adenylate cyclase-activating polypeptide (PACAP),^45^ α-melanocyte-stimulating hormone (α-MSH),^46^ NDA-1^47^). We therefore reasoned that prediction tools for antimicrobial peptides could also aid in the discovery of neuropeptides. By referencing our *de novo* sequences against AMP^48^ and neuropeptide prediction tools,^49,50^ we identified eight peptides with antimicrobial properties, neuropeptide properties, or properties of both classes, including AGLQFPVGR and SSRAGLQFPVGR, which were identified as likely neuropeptides by both prediction models (**Table 1**). One peptide, TLEAKLLR, detected in the PO and CoG (**Fig. 5b, 5c,** **Table 1**), presented characteristics of both AMPs and neuropeptides, with support from both neuropeptide prediction models. Although TLEAKLLR has not been previously reported in the literature, the Database of Antimicrobial Activity and Structure of Peptides (DBAASP)^48^ predicts minor antibacterial activity against *Klebsiella pneumoniae* and antiviral activity against a number of viruses, including SARS-CoV and SARS-CoV-2 (**Supplementary Table 1**). Alignment of these uncharacterized, yet high abundance peptides revealed minimal overlap with known crustacean peptides. AGLQ was the longest consecutive conserved region (**Supplementary Fig. 7**), representing a portion of the hallmark HDAP motif AGLQFPV.^38^ Together, these data identify several candidate bioactive peptides revealed by our hybrid workflow, warranting further functional characterization.

### Discovery of direct homology between invertebrate and vertebrate peptides

Finally, the validated hybrid *de novo*-database search pipeline was applied to investigate endogenous expression due to feeding. Jonah crabs (*Cancer borealis*) are noteworthy model systems for feeding behavior, with reported homology with vertebrate neuropeptides;^2^ however, a direct comparison of crustacean and mammalian feeding-associated peptides has not been reported. We therefore sought to identify novel peptides in these organisms and evaluate cross-species homology. *De novo* peptides from fed Jonah crabs were compared to those from fed rats (*R. norvegicus*). Comparison of these two datasets revealed three pairs of peptides with remarkable homology between the rat peptide and its crustacean counterpart (**Fig. 6**).

**Figure 6:**
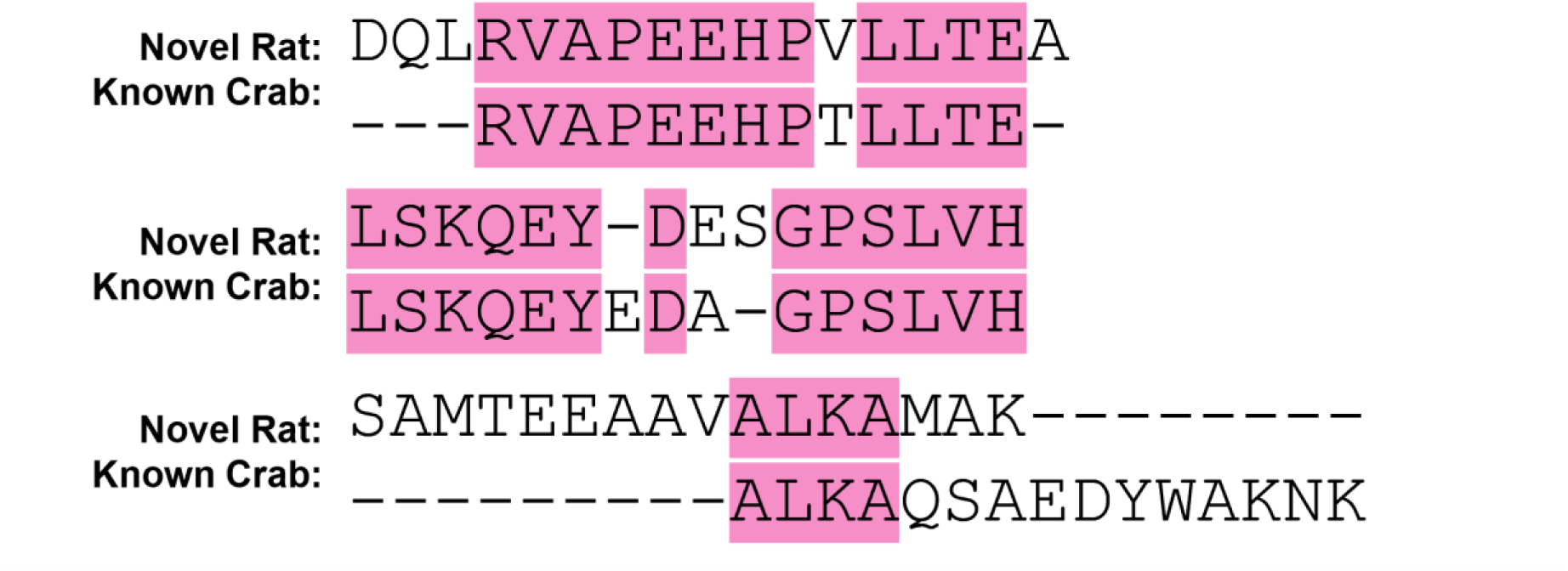
S**e**quence **alignment of novel rat and Jonah crab (C. borealis) peptides**. Novel rat peptides identified by de novo sequencing were aligned with known crustacean peptides to highlight regions of sequence similarity. Conserved residues are highlighted in pink. The conserved regions RVAPEEHP and LSKQEYD correspond to actin, while ALKAMAK corresponds to eukaryotic translation initiation factor 5A-1.

Interestingly, the RVAPEEHP conserved peptide fragment has been defined as expressed by major histocompatibility complex (MHC) I in *Homo sapiens*.^51^ Given that crustaceans lack an adaptive immune system, it is clear that an immunopeptide function would not be induced by this peptide; however, RVAPEEHP as well as LSKQEYD correspond to actin, which is deeply conserved across all animals.^52^ Additionally, ALKAMAK corresponds to a eukaryotic translation initiation factor 5A-1 protein,^53^ suggesting that the aligned ALKA sequence in the crustacean peptide may arise from a crustacean protein with a similar function. Thus, while these conserved peptides do not reflect a signaling peptide or AMP, these findings reflect a new means to annotate housekeeping proteins for sparsely annotated model organisms. Given that the *Cancer borealis* and *Callinectes sapidus* genomes are currently limited to chromosomal-level annotation, it remains difficult to identify peptides that have not been previously characterized through targeted sequencing efforts. Using this hybrid pipeline approach, we can identify high abundance peptides that are broadly expressed, and infer their likely function based on homology with housekeeping proteins in more thoroughly annotated organisms.

## Discussion

In this work, we developed a hybrid analytical workflow integrating EndoGenius, a database search engine designed specifically for endogenous peptides, with Casanovo for de novo peptide sequencing and NovoBoard for empirical FDR control (**Fig. 1**). This pipeline was designed to overcome longstanding challenges in crustacean peptidomics, including incomplete genomes, non-tryptic fragmentation, and the limited compatibility of standard database search tools with non-digested peptides. NovoBoard generated decoy spectra that faithfully captured the statistical properties of experimental data (**Fig. 2a–c**), confirming that FDR estimates derived from these decoys are reliable and independent of database composition. The predicted generalizability of Casanovo was confirmed through generation of well-annotated sequences that retained features characteristic of endogenous, non-digested peptides, including a slight predominance of b-ions (**Figs. 2d, 3a, Supplementary Fig. 5**). Moreover, results at both the de novo and EndoGenius-validated levels reflect the current state of crustacean peptidomics (**Figs. 2e, 3b, 3d**), underscoring the generalizability of Casanovo to this application. While other de novo sequencing platforms exist, Casanovo and NovoBoard are well suited for endogenous peptide analysis, particularly in organisms with limited genomic resources.

Notable variability was observed among the de novo results for technical replicates of each tissue (**Fig. 2e, Supplementary Fig. 4**). This is likely a consequence of the stochastic nature of DDA, which inherently biases against low-abundance analytes such as neuropeptides and other endogenous peptides.^2^ Data-independent acquisition (DIA) has been applied to overcome this bias by fragmenting all ions within a given *m/z* window rather than only the *n*-most abundant ions,^54^ and has even shown to be effective for crustacean neuropeptides.^7,55^ The manner in which DIA acquires data results in complex, often chimeric spectra, with peptide fragment ions distributed across multiple scans, complicating both identification and de novo sequencing. Analytical strategies have been developed for *de novo* sequencing of DIA data,^20,56–58^ which have been successfully applied to proteomics datasets. This establishes a promising avenue of future investigation for analysis of crustacean peptidomics and other non-digested peptide datasets.

Validated *de novo* peptides from the blue crab dataset revealed novel peptides that were unidentifiable by other search techniques (**Figs. 3c, 3d, Supplementary Fig. 6**). A global alignment of these peptides with known crustacean neuropeptides as well as those from other organisms revealed marked homology (**Fig. 3e**), supporting the putative classification of some of these peptides as neuropeptides. Additional homology-based analyses revealed that AGLQFPVGR, the only peptide shared by all tissues, was identified as a histone-directed protein based on alignment with related sequences in invertebrates including several crustacean and insect species,^34–37^ as well as vertebrates including fish and mammals.^33,36^ Studies suggest that this peptide and others like it play a role in host defense, particularly for animals such as crustaceans that lack an adaptive immune system, leading to its characterization as a histone-derived antimicrobial peptide (HDAP). This is the first report of this HDAP in the blue crab, which we find to be broadly expressed across several chromosomes within the *C. sapidus* genome (**Fig. 4a**).

Notably, the encoding gene is broadly expressed across several chromosomes within the C. sapidus genome. Only one chromosome retains the exact sequence of interest, whereas six other chromosomes all exhibit the same four amino acid substitutions. Structural predictions reveal that the substituted residues co-localize in space (**Fig. 4c**), suggesting potential differences in binding affinity or other physiochemical properties, despite conservation of the overall helical fold. Further investigations is needed to determine the functional role of AGLQFPVGR in blue crabs, as homology alone does not conclusively establish function. Interestingly, neither AGLQFPVGR nor SSRAGLQFPVGR were predicted to have antimicrobial activity according to DBAASP (**Table 1**), raising the possibility that this peptide may serve a different function in blue crabs than other species. The variant forms also warrant investigation, given their genomic prevalence and the potential for altered physiochemical properties. Ultimately, the identification and characterization of a widely conserved HDAP-derived peptide in C. sapidus demonstrates the capacity of this hybrid approach to yield biologically meaningful discoveries.

Novel peptides demonstrated a wide dynamic range across tissue types (**Fig. 5**), with several identified as potential AMPs or neuropeptides using activity prediction tools.^48–50^ Only one peptide was predicted to have both antimicrobial and neuropeptide activity, TLEAKLLR (**Table 1**), with a number of antiviral predictions (**Supplementary Table 1**). This peptide, among others described in **Table 1**, are under-characterized or wholly uncharacterized, and therefore represent promising targets for functional characterization, particularly regarding their potential roles in innate immunity and neuromodulation in *C. sapidus*.

Lastly, we were able to extend our validated workflow to offer the first direct comparison of crustacean and mammalian peptides expressed during feeding. This was not previously achievable due to the incompatibility in annotation of crustacean and mammalian (*e.g.*, rats) genomes. *De novo* sequencing via our hybrid workflow enabled identification of crustacean peptides not present in existing databases, thereby enabling direct comparison with mammalian peptides. With this hybrid workflow, we identified three peptides from Jonah crabs (*C. borealis*) with notable homology with rats, each of which is related to housekeeping proteins in its mammalian counterpart. Although these peptides do not appear to be neuropeptides or AMPs, they represent the first characterization of housekeeping proteins in Jonah crabs. Future work applying *de novo* sequencing and hybrid pipelines to improve genome annotation has the potential to provide a more comprehensive understanding of the proteomic as well as peptidomic framework for the critical model organisms.

Together, this work presents a hybrid peptide discovery platform which combines transformer-based de novo sequencing with EndoGenius, a platform optimized for endogenous peptides. This framework, integrating Casanovo, EndoGenius, and FDR control through NovoBoard, overcomes longstanding limitations in primary research using crustacean model systems, including incomplete genomes, non-tryptic fragmentation, and the poor detectability of modified or degraded peptides, while substantially expanding the accessible sequence space. Most notably, we report the first histone-2A-derived AMP in the blue crab, define its genomic organization, and characterize naturally occurring sequence variants. We also reveal several high-abundance novel neuropeptides across neuroendocrine tissues, including peptides with predicted antimicrobial or metabolic regulatory activity, and uncover previously unrecognized cross-phyla peptide homologies which improve annotation of housekeeping proteins in crustaceans. These findings demonstrate that our hybrid platform increases the confidence and depth of peptide identifications, while also enabling biologically significant discoveries that were previously inaccessible. This highlights the transformative potential of combining modern *de novo* sequencing with tailored bioinformatics to accelerate peptide discovery in emerging model systems.

## Methods

### Chemicals and Materials

Unless otherwise noted, all reagents were Optima grade and purchased from Fisher Scientific (Pittsburgh, PA). Formic acid was purchased from Sigma-Aldrich (St. Louis, MO). Notable solutions in this work include methanol (MeOH) acidified with acetic acid (AA), prepared as 90/9/1 (v/v/v) for MeOH/H_2_O/AA, and physiological saline, prepared as 440 mM NaCl, 11 mM KCl, 13 mM CaCl_2_, 26 mM MgCl_2_, 10 mM Trizma base, and 5 mM maleic acid, adjusted to pH 7.4 with NaOH.

### Animals and Feeding Experiments

Initial evaluation of the platform was conducted with female blue crabs (*Callinectes sapidus*), while the feeding study was conducted with male Jonah crabs (*Cancer borealis*). Both species of crabs were purchased from Global Market in Madison, WI and transferred to an artificial seawater environment composed of 35 parts per thousand (ppt) salinity, 13-16 °C, and 8-10 ppm O_2_. All animals were allowed to acclimate to their environment for at least two weeks prior to any experimentation. Although no institutional approval is required for housing, caring for, and otherwise using crustaceans in experiments, all experiments were performed in accordance with local and national regulations.

Crabs were anesthetized on ice for 30 min prior to sacrifice. The brain, paired sinus glands (SG), thoracic ganglion (TG), paired commissural ganglia (CoG), and paired pericardial organs (PO) were collected, as previously described.^59^ Dissections were conducted in chilled crab physiological saline, and tissues were heat stabilized (Denator Stabilizer T1, Gothenburg, Sweden) to halt endogenous protease activity, after which they were flash frozen on dry ice.

For crustacean feeding experiments, Jonah crabs were each fed 4.0 g tilapia, achieving satiety. After crabs finished consuming the food (∼30 min), an additional 30 min was allowed to elapse before crabs were anesthetized and euthanized as described above. Data from fed adult male Sprague-Dawley rats (*Rattus norvegicus*) were reanalyzed from a previous study.^60^

### Peptide Extraction and Desalting

Tissues were submerged in 200 μL chilled acidified MeOH and homogenized using a probe ultrasonication device (Fisher Scientific, Pittsburgh, PA). Fully homogenized samples were centrifuged, after which the supernatant was collected and dried via vacuum centrifuge. Further details regarding the extraction process have been described previously.^61^ Dried samples were reconstituted and biological replicates (*n* = 3) were pooled for each tissue type. Pooled samples were desalted via P100 C18 OMIX pipette tips (Agilent, Santa Clara, CA) according to the manufacturer’s instructions.

### Mass Spectrometry Data Acquisition

Data for evaluation of the hybrid pipeline were acquired from *C. sapidus* tissue using a Thermo Q-Exactive HF mass spectrometer coupled to a Dionex Ultimate 3000 LC system (Thermo Scientific, Waltham, MA). For the feeding study, data were acquired from *C. borealis* tissue using a Thermo Q-Exactive mass spectrometer coupled to a Waters nanoAcquity LC system (Waters Corp, Milford, MA, USA). Instrument method parameters for both acquisitions were kept consistent with previous studies.^26^ Identical chromatographic conditions were used for both datasets.

Chromatographic separation was performed on an in-house prepared 15 cm microcapillary column with an integrated emitter tip, packed with 1.7 μm bridged ethylene hybrid C18 particles. Mobile phases A and B were Optima-grade water and acetonitrile, respectively, each with 0.1% (v/v) formic acid (FA). Separation was carried out at 0.300 μL/min using a 120 min gradient as follows (% mobile phase B): 0–1 min, 3–10%; 1–90 min, 10–35%; 90–92 min, 35–95%; 92–102 min, 95%; 102–105 min, 95–3%; 105–120 min, 3%. Samples were reconstituted in 20 μL 0.1% FA in water, and 2 μL were injected for each of three technical replicate analyses per sample.

Data were acquired in data-dependent acquisition (DDA) mode, acquiring the top 20 precursors for MS^2^ acquisition under positive electrospray ionization (ESI). Pertinent parameter settings included collision energy, 30 eV; MS^1^ resolution, 60k; MS^2^ resolution, 15k; isolation window, 2.0 *m/z*; dynamic exclusion, 50 s, and scan range, 200-2000 *m/z*.

### Software Notes

Database searching via EndoGenius (v1.1.4)^26,62^ and post-analysis processing were performed on a workstation equipped with a 64-bit quad-core processor (3.10 GHz) and 128 GB RAM running Windows 10. De novo sequencing was conducted using Casanovo (v4.2.0; Docker image casanovo:4.2.1)¹⁴’²³ and FDR estimation via NovoBoard²⁴ on the Center for High Throughput Computing (CHTC) at the University of Wisconsin-Madison,^63^ utilizing CPU-based high throughput resources, with typical requests of 6 CPUs, 32 GB memory, and 40 GB disk per task. Raw file conversion was performed using MSConvert (v3.0)⁶² and RawConverter (v1.1.0.18).⁶³ Predicted structures were generated using the AlphaFold Server (https://alphafoldserver.com), powered by AlphaFold 3,³²’³³ and visualized in PyMOL (v3.1.4.1).³⁴

Antimicrobial peptide activity was predicted using DBAASP⁴¹ and neuropeptide activity was assessed using NeuroPred-FRL⁴² and NeuroPP.⁴³ All custom scripts used for processing *de novo* sequencing results are available at https://github.com/lingjunli-research/HybridEndoGenius.

### Data Analysis via EndoGenius

Raw files were converted to mzML and MS2 formats using MSConvert^64^ and RawConverter,^65^ respectively, using default settings for each. Data were searched in EndoGenius against an in-house developed database containing 891 peptides concatenated with decoy sequences. Error thresholds for precursor and fragment masses were set to 20 ppm and 0.02 Da, respectively. Variable modifications included C-terminal amidation, N-terminal cyclization of glutamine and glutamic acid (pyro-glu), and oxidation on methionine, with up to three modifications allowed per peptide. Other relevant parameters include a minimum m/z value of 50 and a minimum intensity of 1000. Search results were filtered at an EndoGenius confidence score of 1000, which corresponding to an estimated FDR of less than 5%.^26^

### De Novo Sequencing

In addition to the mzML files generated previously, raw files were also converted to MGF format using MSConvert.^66^ Both formats were used to ensure consistent peak representation while retaining spectrum-level tracking information.^67^

Paired decoy spectra were generated from each target MS/MS spectrum using NovoBoard with a 50% peak-removal setting, as previously benchmarked for Casanovo.²⁴ Target and decoy MGF files were processed with Casanovo (v4.2.0; Docker image casanovo:4.2.1)¹⁴’²³ using default configuration parameters to generate de novo sequence predictions and associated confidence scores for every peptide-spectrum match (PSM). Confidence scores were returned to NovoBoard to estimate per-file FDR thresholds, and a 5% FDR cutoff was applied.²⁴ Filtered identifications were compiled into FASTA format and used as input for a second round of database searching via EndoGenius using the same parameters as described above.

### Post-analysis manipulation and calculations

Predicted structures for the peptide AGLQFPVGR were generated using the AlphaFold Server^39,40^ with default parameters and visualized using PyMol.^41^ Novel peptide sequences were assessed for antimicrobial activity via the Database of Antimicrobial Activity and Structure of Peptides (DBAASP),^48^ and for neuropeptide activity via NeuroPred-FRL^49^ and NeuroPP.^50^ Sequences were submitted to each tool in FASTA format using default parameters.

Peptide coverage by b- and y-ions was calculated according to Equations 1 and 2, where *b* and *y* respectively represent the number of standard b- and y-ions identified, b_neut_ and y_neut_ respectively represent the number of identified b- and y-ions considering a neutral loss (*e.g.*, loss of H_2_O or NH_3_), and b_theory_ and y_theory_ respectively represent the number theoretically available of b- and y-ions.

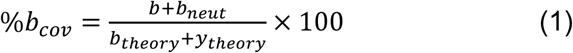

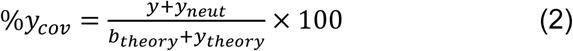

### Statistical analyses

All analyses were performed using three technical replicate injections per tissue. Peptide abundance was quantified as log-transformed precursor intensity, and error bars and shading in Figures 3b and 5 represent the standard deviation across technical replicates. Kernel density estimates for target and decoy spectral feature distributions (**Figs. 2a–c, Supplementary Figs. 1–3**) were generated using Python with default bandwidth parameters. Peptide identification overlap across replicates and search strategies was assessed using Venn diagrams representing unique and shared identifications. Sequence similarity between novel de novo peptides and known neuropeptide database entries (**Fig. 3e**) was calculated as the percentage of aligned residues using Clustal Omega. Rank-abundance profiles (**Fig. 5**) were generated by ranking all identified peptides by mean precursor intensity across technical replicates. False discovery rate control was applied at 5% for both database searching (EndoGenius confidence score of 1000) and de novo sequencing (NovoBoard per-file score threshold).

## Data availability

Mass spectrometry peptidomic data for the crustacean datasets generated and analyzed here as well as the rat data dataset generated previously^60^ and analyzed here were deposited to the ProteomeXchange Consortium via the MassIVE partner repository. The identifiers are MSV000101523 and MSV000080106 for the crustacean and rat datasets, respectively.

## Code Availability

Code regarding use of EndoGenius can be found at https://github.com/lingjunli-research/EndoGenius. Additional code used for processing *de novo* sequencing results are included at this location. For increased accessibility, a graphical user interface is provided for EndoGenius usage.

## Acknowledgements

Computational analyses, including Casanovo and EndoGenius, were performed at the Center for High Throughput Computing (CHTC) at the University of Wisconsin- Madison.^68^ This work was supported in part by National Institutes of Health (NIH) through grants R01DK071801 and R01NS029346 (L.L.) and the National Science Foundation (NSF) through the grant CHE-2108223 (L.L.).This work was supported in part by National Institutes of Health (NIH) through grants R01AG078794 (L.L.), R01DK071801 (L.L.), and the Research Forward grant by University of Wisconsin - Madison Office of the Vice Chancellor for Research with funding from the Wisconsin Alumni Research Foundation (L.L.). The Orbitrap instruments were purchased through the support of an NIH shared instrument grant S10RR029531 and Office of the Vice Chancellor for Research and Graduate Education at the University of Wisconsin-Madison. L.F. was supported in part by the National Institute of General Medical Sciences of the National Institutes of Health under Award Number T32GM008505 (Chemistry–Biology Interface Training Program), the 2024 Eli Lilly and Company/ACS Analytical Graduate Fellowship, and a predoctoral fellowship supported by the NIH, under Ruth L. Kirschstein National Research Service Award (NRSA) from the National Institutes of Health-General Medical Sciences F31GM156104. A.E.I was supported in part by the National Science Foundation Graduate Research Fellowship Program (GRFP) under Grant No. 2137424. Any opinions, findings, and conclusions or recommendations expressed in this material are those of the author(s) and do not necessarily reflect the views of the National Science Foundation. T.C.D was supported in part by the National Institute of General Medical Services of the National Institute of Health under Award 5T32GM141013 (Molecular and Cellular Pharmacology Training Program) and a SciMed Graduate Research Scholars Fellowship through the University of Wisconsin-Madison. L.L. would like to acknowledge NIH grants R01AG052324, R01AG078794, S10OD028473, and S10OD025084, as well as funding support from a Vilas Distinguished Achievement Professorship and Charles Melbourne Johnson Professorship with funding provided by the Wisconsin Alumni Research Foundation and University of Wisconsin-Madison School of Pharmacy. Figure 1 was generated with Biorender.

## Author information

### Authors and Affiliations

Department of Chemistry, University of Wisconsin-Madison, Madison, Wisconsin, 53706, USA

Lauren Fields, Angel E. Ibarra, Kendra G. Selby, Tong Gao, Lingjun Li.

Department of Statistics, University of Wisconsin-Madison, 1205 University Avenue, Madison, WI, 53706, USA

Jiangrong Qin

Department of Mathematics, University of Wisconsin-Madison, 480 Lincoln Drive, Madison, WI, 53706, USA

Jiangrong Qin

School of Pharmacy, University of Wisconsin-Madison, Madison, Wisconsin, 53705, USA

Tina C. Dang, Haiyan Lu, Lingjun Li

Lachman Institute for Pharmaceutical Development, School of Pharmacy, University of Wisconsin-Madison, Madison, WI, 53705, USA

Lingjun Li

Wisconsin Center for NanoBioSystems, School of Pharmacy, University of Wisconsin-Madison, Madison, WI, 53705, USA

Lingjun Li

### Contributions

L.F. and L.L. designed the research; L.F. collected tissues and prepared samples for MS acquisition. J.Q. developed computational pipeline and performed cluster analyses. L.F., J.Q., A.E.I., K.G.S., T.G., T.C.D., H.L., and L.L analyzed data. L.F., J.Q., and L.L. prepared the manuscript. K.G.S. and T.G. assisted in figure generation. All authors contributed to the discussion of the project and have reviewed and approved of the final manuscript.

*Corresponding author* Lingjun Li

---

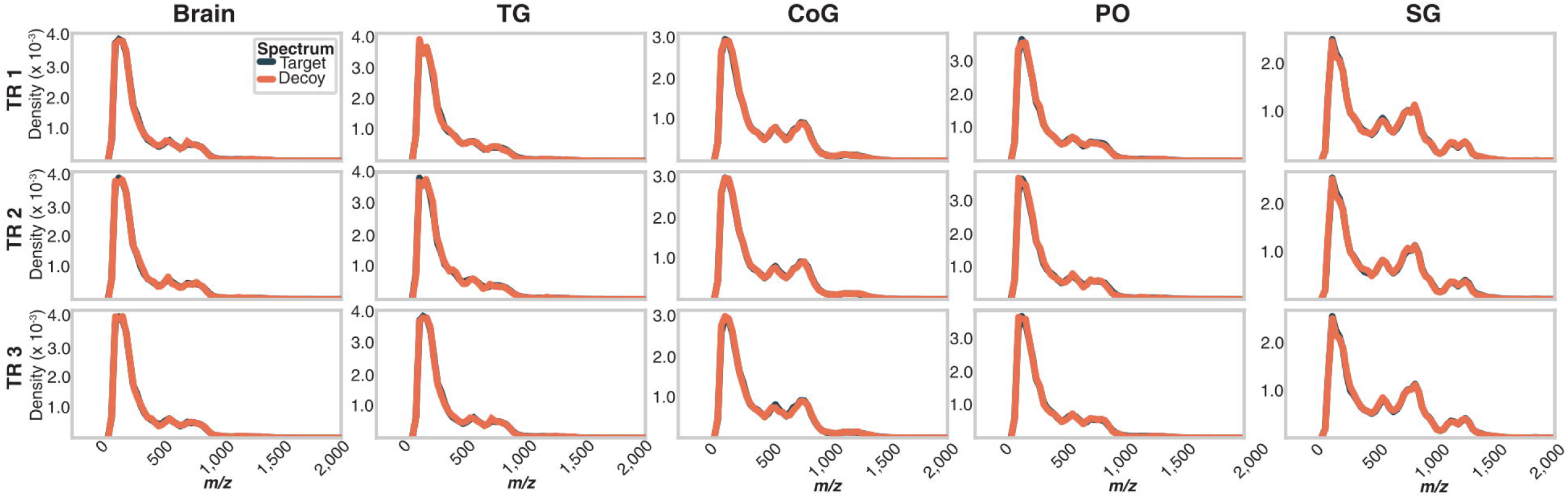

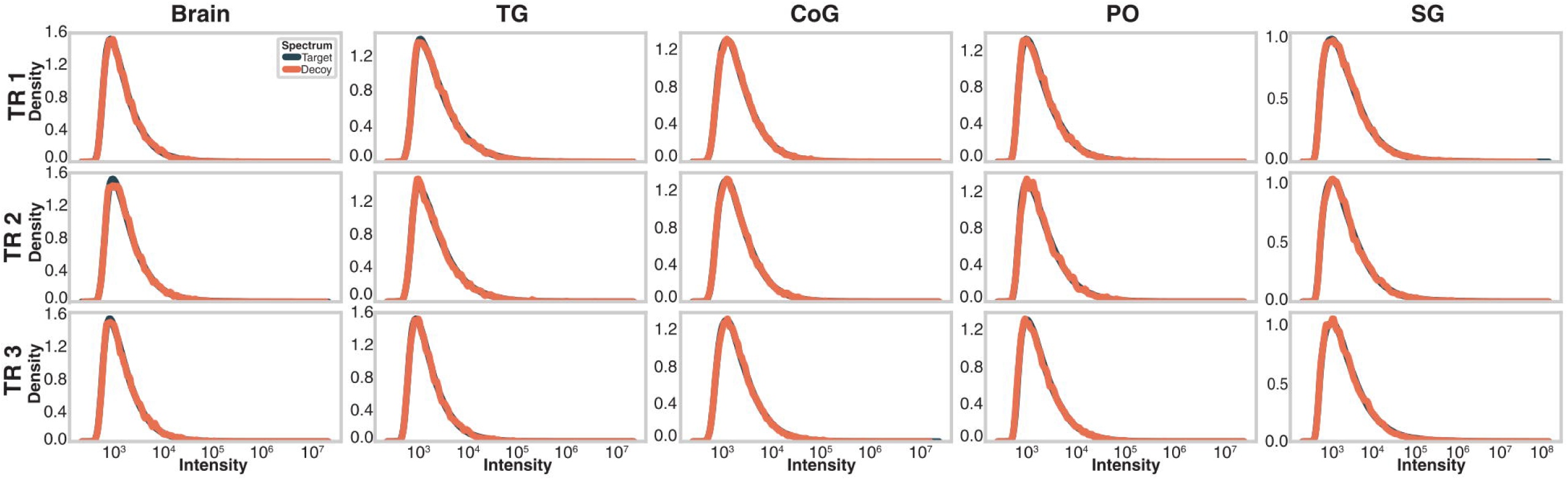

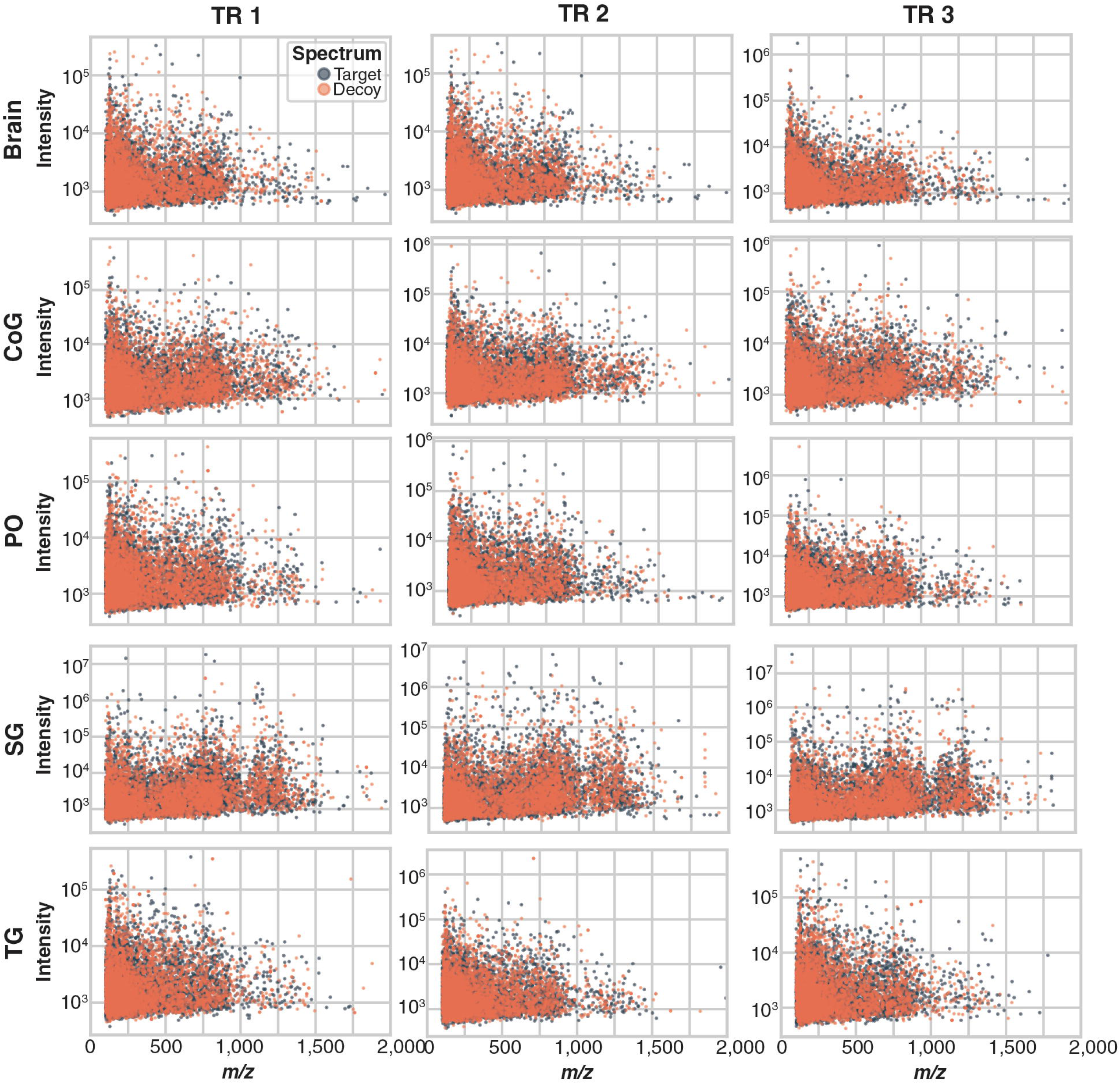

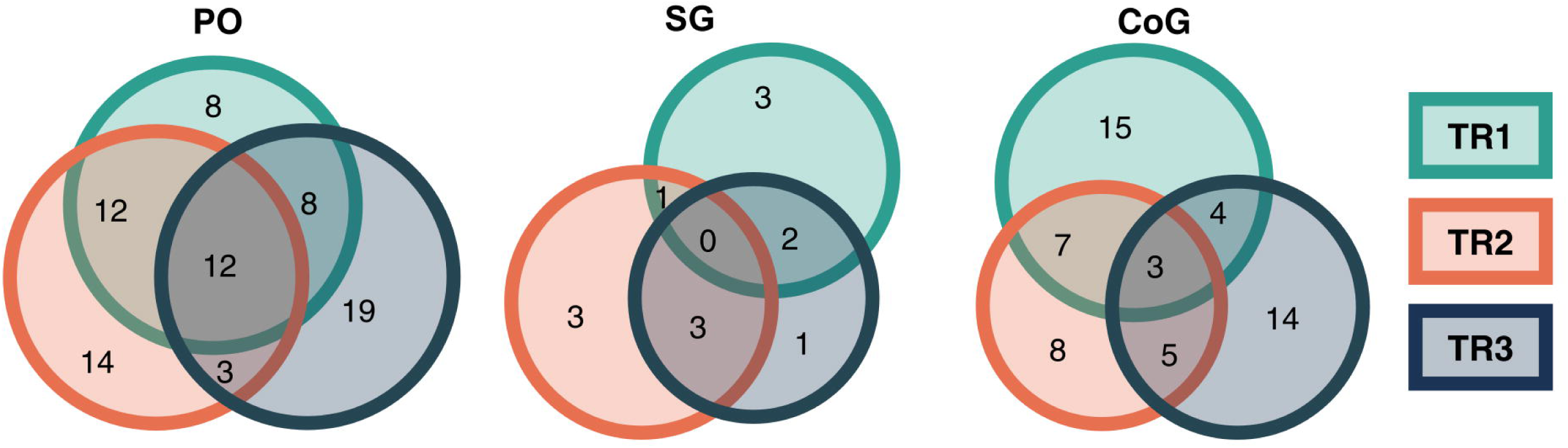

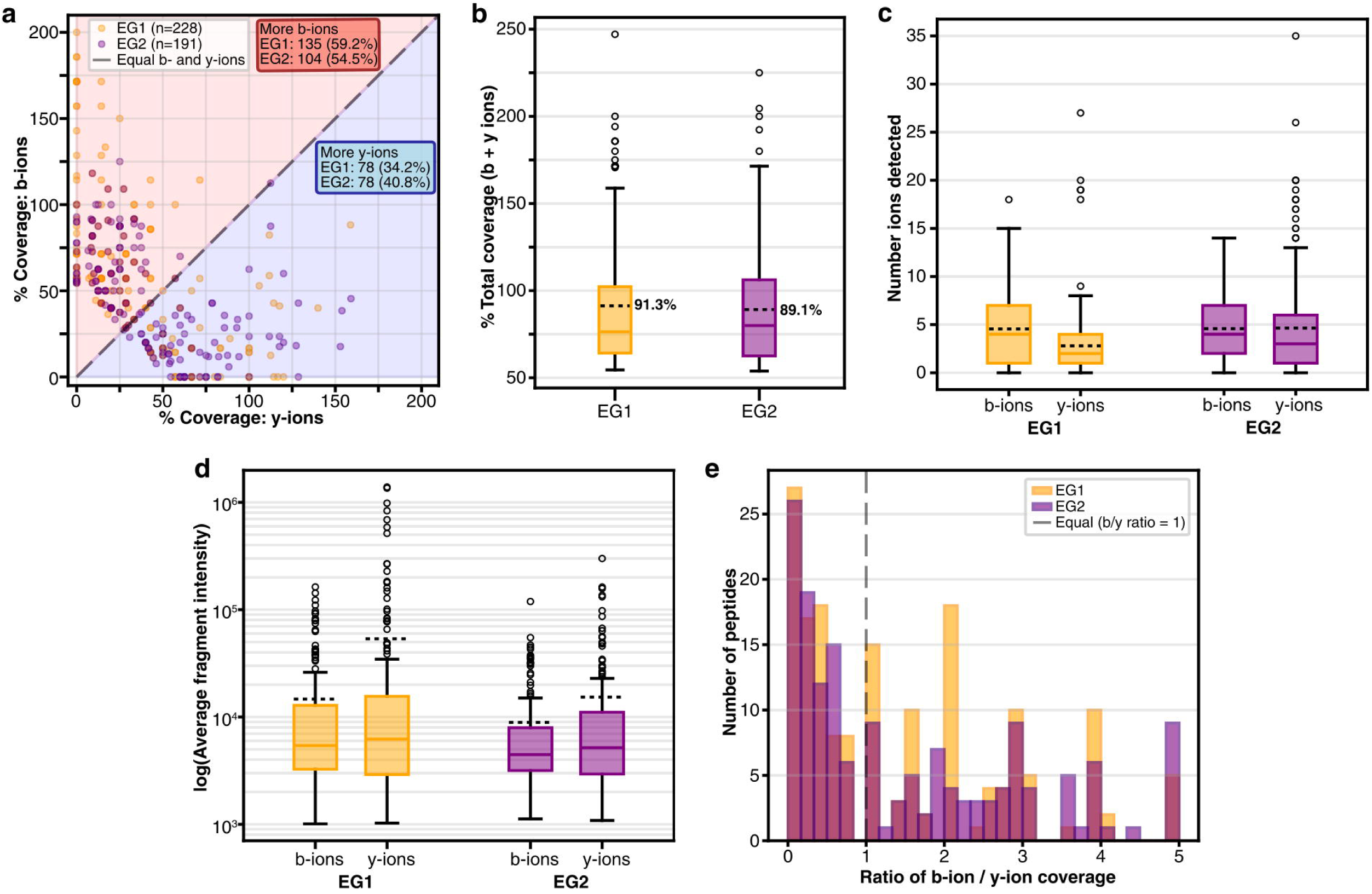

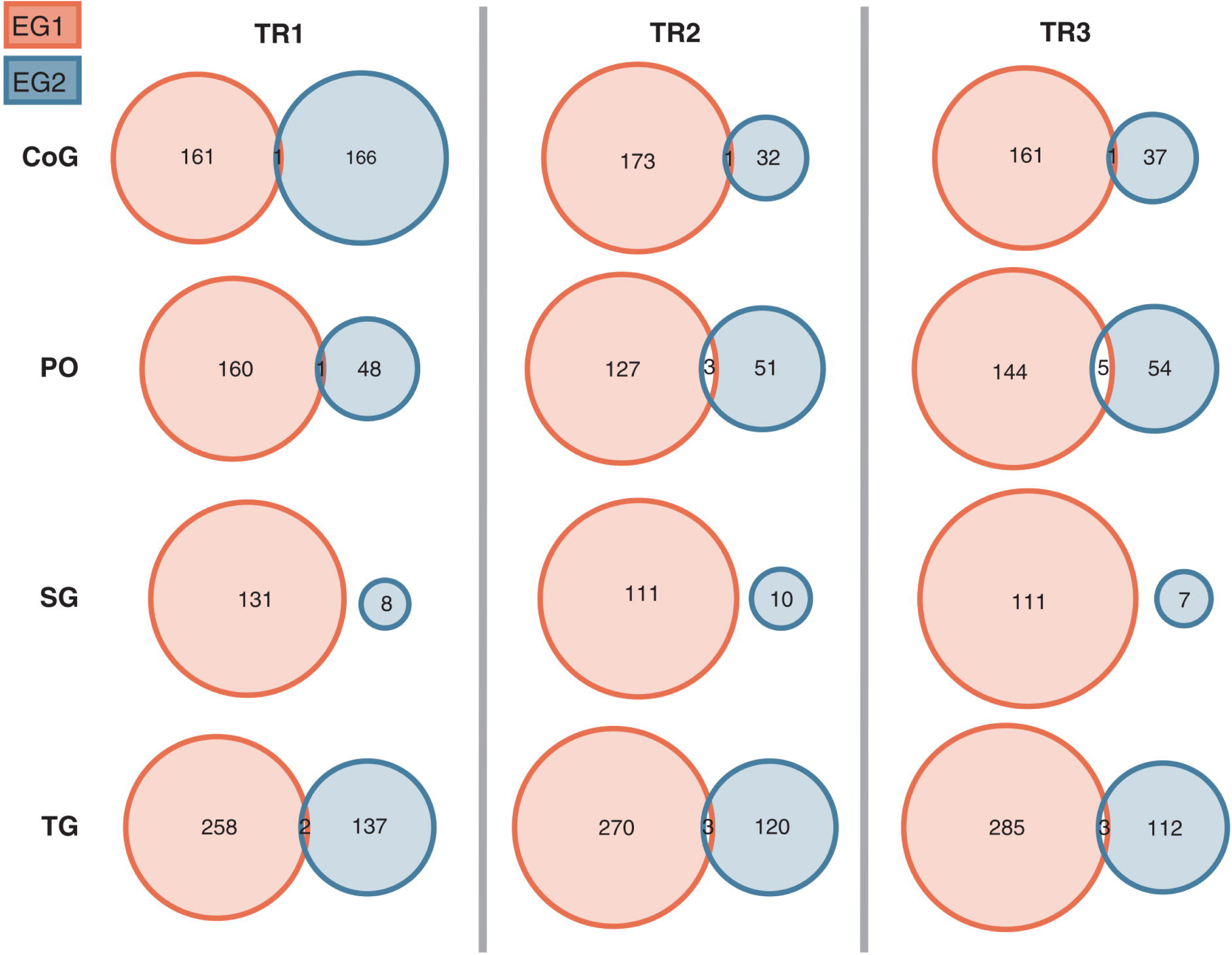

