## Supplemental Information for "Bridging genomes and peptidomes: hybrid sequencing reveals conserved bioactive peptides in crustaceans"

**Table of Contents**

Supplemental Figures and Tables (located within this document)


- **Figure S1:** Tissue-specific fragment *m/z* density for target and NovoBoard-generated decoy sequences.
- **Figure S2:** Tissue-specific fragment intensity density for target and NovoBoard-generated decoy sequences.
- **Figure S3.** Tissue-specific fragment intensity vs *m/z* for target and NovoBoard-generated decoy sequences.
- **Figure S4:** Overlap in *de novo* sequencing results for PO, SG, and CoG.
- **Figure S5:** Comparison of b- and y-ion coverage by EndoGenius search results.
- **Figure S6:** Overlap in identifications for EndoGenius analyses of spectra directly and *de novo* results.
- **Figure S7:** Top nine novel peptides aligned with the most similar peptide from the crustacean neuropeptide database.
- **Table S1:** Prediction of antibacterial, antifungal, and antiviral activity for peptide TLEAKLLR using the Database of Antimicrobial Activity and Structure of Peptides (DBAASP).
- **Table S2:** Alignment of histone-derived antimicrobial peptide (HDAP) across crustacea.

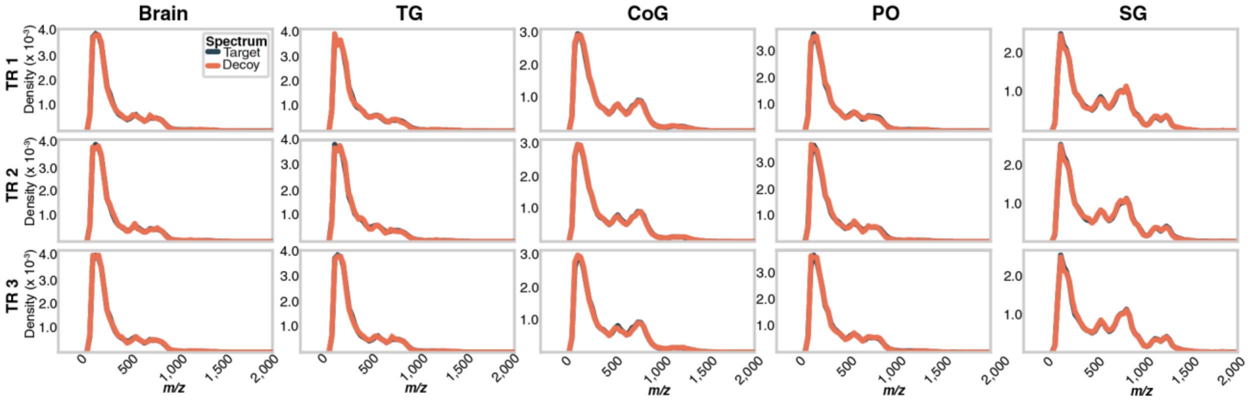


**Supplementary Figure 1. Tissue-specific fragment *m/z* density for target and NovoBoard-generated decoy sequences.** Traces are shown for all crustacean neuroendocrine tissues for each of three technical replicates (TR). Target (dark teal) and decoy (coral) traces overlap closely in all panels, indicating strong agreement between target and decoy spectral characteristics. Abbreviations: thoracic ganglion (TG), commissural ganglia (CoG), pericardial organs (PO), and sinus glands (SG).

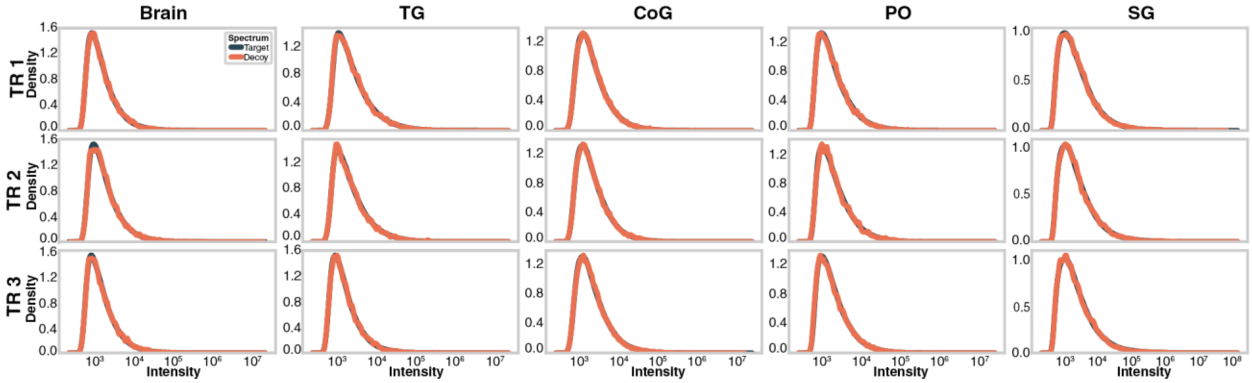


**Supplementary Figure 2. Tissue-specific fragment intensity density for target and NovoBoard-generated decoy sequences.** Traces are shown all crustacean neuroendocrine tissues for each of three technical replicates (TR). Target (dark teal) and decoy (coral) traces overlap closely in all panels, indicating strong agreement between target and decoy spectral characteristics. Abbreviations: thoracic ganglion (TG), commissural ganglia (CoG), pericardial organs (PO), and sinus glands (SG).

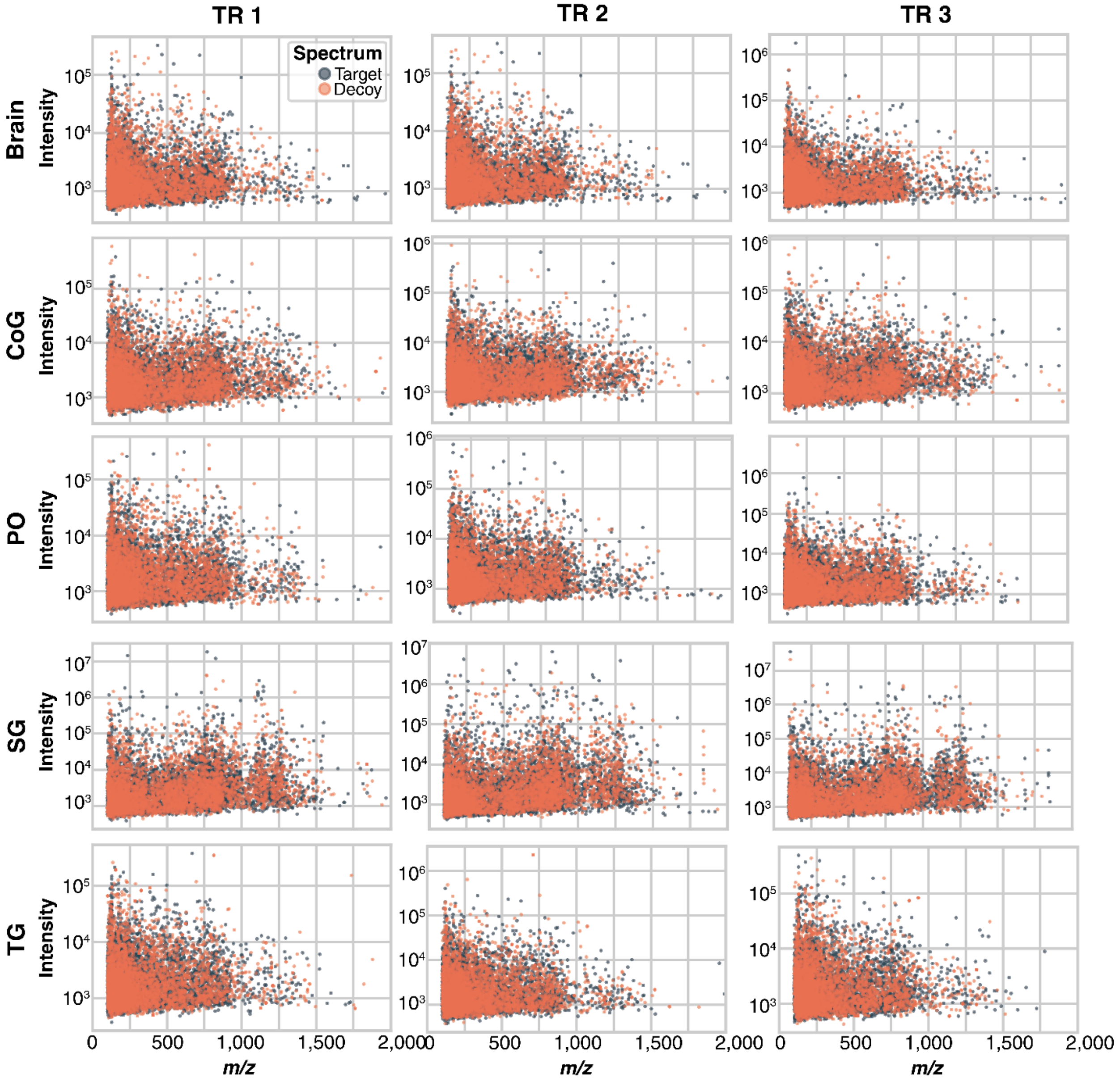


**Supplementary Figure 3. Tissue-specific fragment intensity vs *m/z* for target and NovoBoard-generated decoy sequences.** Each panel shows the joint distribution of fragment m/z and intensity for all spectra in a given tissue and technical replicate. Abbreviations: thoracic ganglion (TG), commissural ganglia (CoG), pericardial organs (PO), and sinus glands (SG).

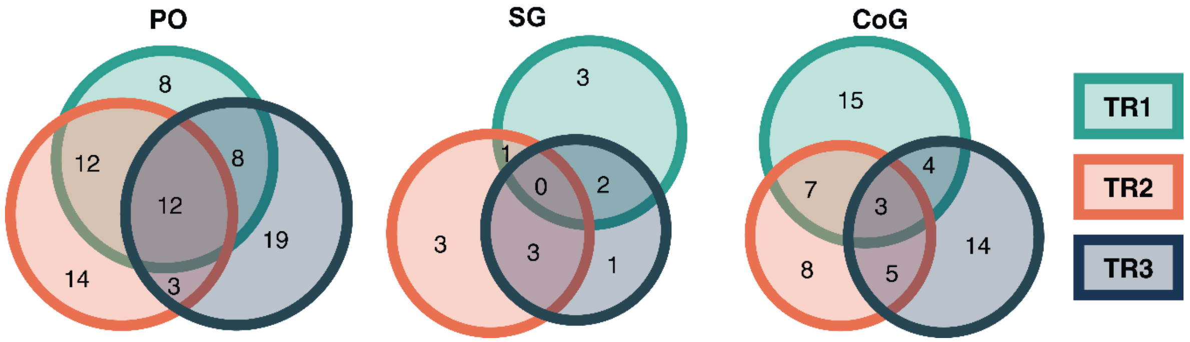


**Supplementary Figure 4. Overlap in *de novo* sequencing results for PO, SG, and CoG.** Three technical replicate injections (TR) were collected for each sample. Note that no peptides were shared across all three replicates for the SG. Abbreviations: pericardial organs (PO), sinus glands (SG), and commissural ganglia (CoG).

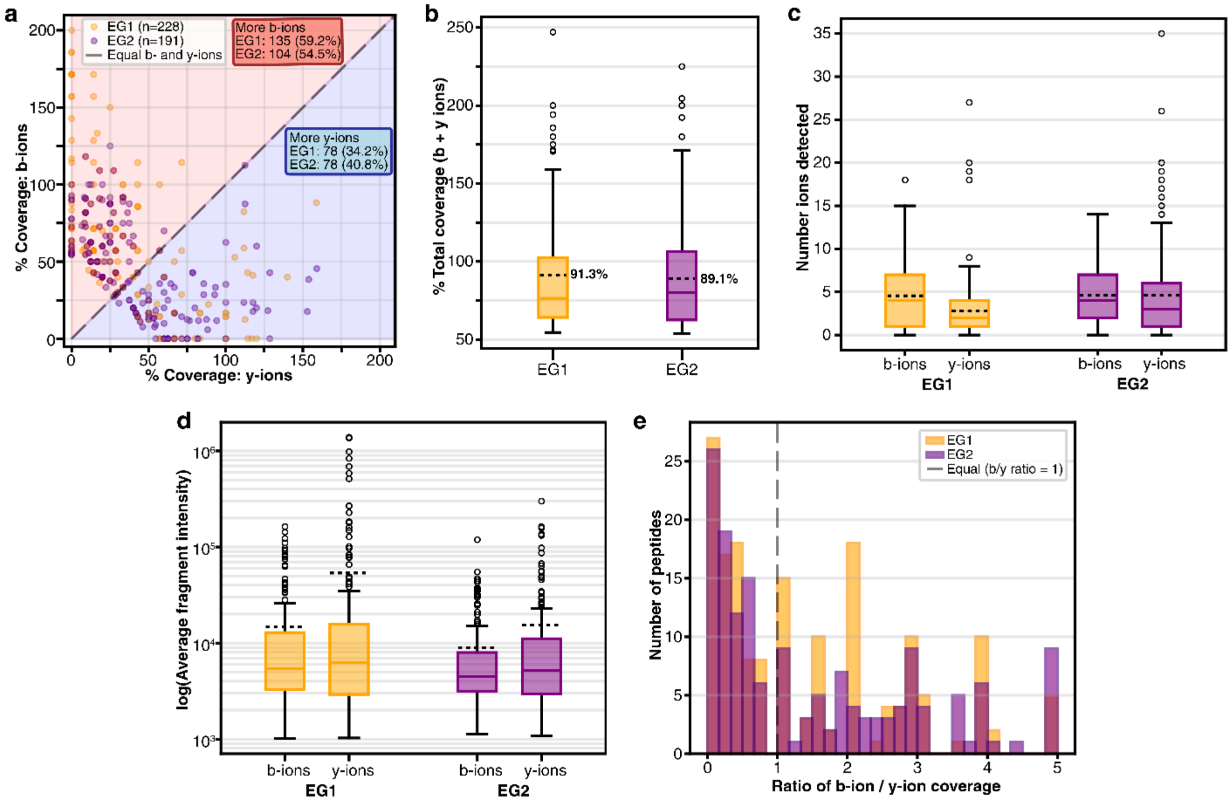


**Supplementary Figure 5. Comparison of b- and y-ion coverage by EndoGenius search results.** EG1 refers to the direct EndoGenius database search of MS spectra, whereas EG2 refers to the combined results of the direct search and the EndoGenius validation search of

Casanovo de novo predictions. **a)** b- versus y-ion coverage for individual peptides, **b)** total ion coverage (b + y), **c)** number of b- and y-ions detected per peptide, **d)** average fragment intensity for b- and y-ions, and **e)** distribution of b-ion to y-ion coverage ratios before (EG1) and after (EG2) integration of de novo sequencing results.

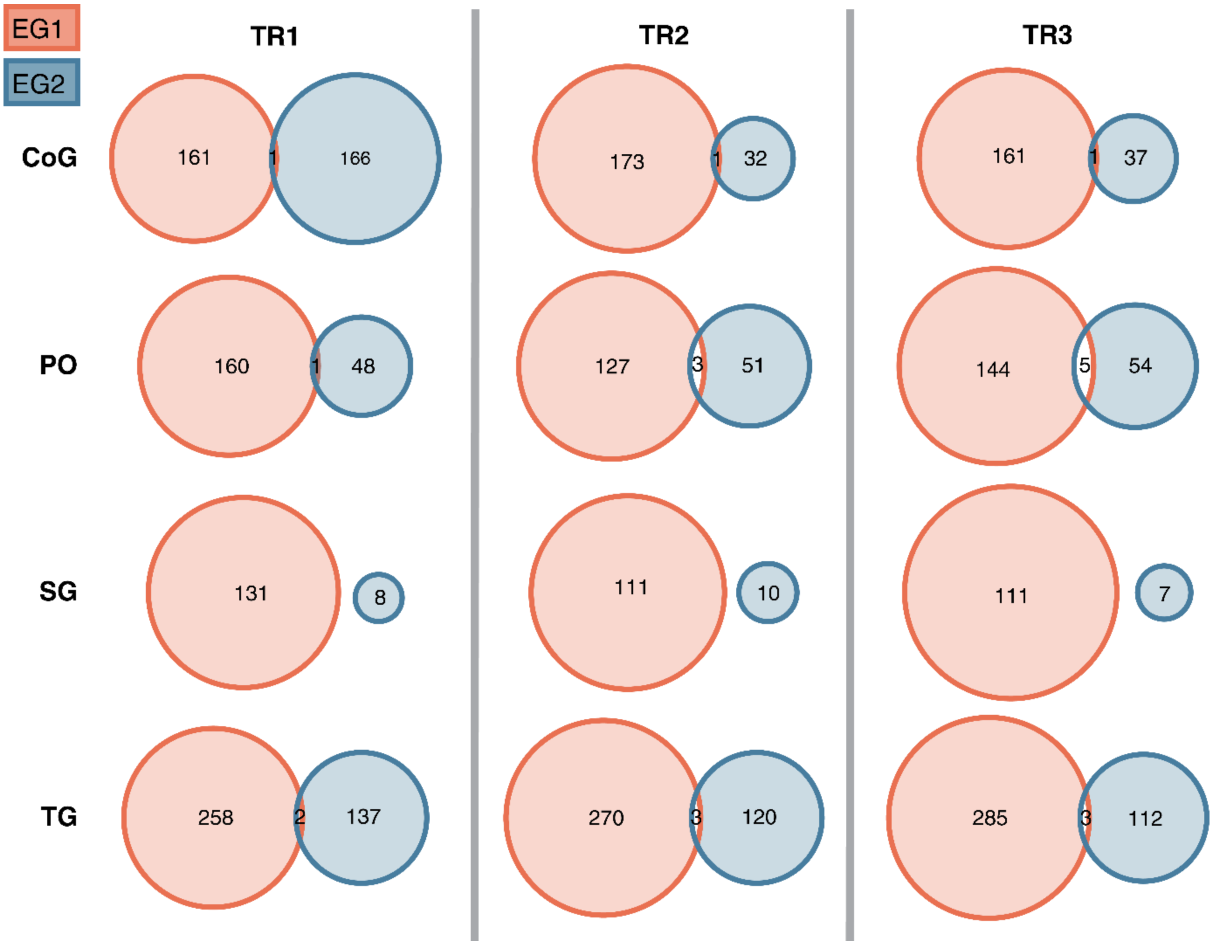


**Supplementary Figure 6. Overlap between direct database search (EG1) and de novo-validated (EG2) identifications across tissues and technical replicates.** EG1 (coral) represents direct EndoGenius database search results and EG2 (blue) represents EndoGenius-validated de novo predictions. Results are shown for each tissue across three technical replicates (TR). Abbreviations: thoracic ganglion (TG), commissural ganglia (CoG), pericardial organs (PO), and sinus glands (SG).

| | | | |
|---|---|---|---|
| **Novel** | S--TLTSRELQTAVR | GS-FMSVESLSR- | VVQWGG-R- |
| **Known** | SMPTLRLRF------ | -SEF--VFS-SRP | FV--GGSRY |
| **Novel** | SSRAGLQFPVG--R- | SNTTALAEAWAR | TLEAKLLR- |
| **Known** | SSR----F-VGGSRY | ----PSFNAWA- | -LELNFLRF |
| **Novel** | -AKWDAWNS-R--- | -AGLQFPVGR | FFVHWNR |
| **Known** | SA--D-WNSLRGTW | AAGLQNYDFG | -FVNSRY |

**Supplementary Figure 7. Top nine novel peptides aligned with the most similar peptide from the in-house crustacean neuropeptide database.** ‘Novel’ refers to peptides identified by de novo sequencing, and 'Known' refers to the most similar sequence in the in-house crustacean neuropeptide database.

**Table S1. Prediction of antibacterial, antifungal, and antiviral activity for peptide TLEAKLLR using the Database of Antimicrobial Activity and Structure of Peptides (DBAASP).**

| Antibacterial prediction[a] | | | | |
|---|---|---|---|---|
| | Machine Learning Approach | | Clusterization Approach | |
| | Activity Prediction | NPV[b] | Activity Prediction | NPV or PPV[b] |
| *Escherichia coli* | Not active | 0.88 | Not Active | 0.84 (NPV) |
| *Pseudomonas aeruginosa* | Not active | 0.67 | Not Active | 0.68 (NPV) |
| *Klebsiella pneumoniae* | Not active | 0.77 | Active | 0.90 (PPV) |
| *Staphylococcus aureus* | Not active | 0.82 | Not Active | 0.80 (NPV) |
| *Bacillus subtilis* | Not active | 0.88 | Not Active | 0.76 (NPV) |
| **Antifungal prediction** | | | | |
| | Activity Prediction | | NPV | |
| *Candida albicans* | Not active | | 0.7 | |
| *Saccharomyces cerevisiae* | Not active | | 0.7 | |
| **Antiviral prediction** | | | | |
| | Activity Prediction | | NPV | |
| DENV-1 | Active | | 0.71 | |
| DENV-2 | Active | | 0.65 | |
| HIV-1 | Active | | 0.57 | |
| Japanese encephalitis virus (JEV) | Active | | 0.87 | |
| MERS-CoV | Active | | 0.61 | |
| SARS-CoV | Active | | 0.64 | |
| SARS-CoV-2 | Active | | 0.61 | |
| West Nile virus (WNV) | Active | | 0.59 | |
| Zika virus (ZIKV) | Active | | 0.66 | |
| Hepatitis C virus (HCV) | Active | | 0.9 | |
| HSV-1 | Not Active | | 0.6 | |
| Vesicular Stomatitis Virus (VSV) | Not Active | | 0.65 | |

[a]Both antibacterial prediction tools also reference peptide sequence data
[b]Negative predictive value (NPV) and positive predictive value (PPV).

**Table S2.** Alignment of histone-derived antimicrobial peptide (HDAP) across crustacea.

| Species | Chr | Seq |
|---|---|---|
| Query | - | RSSRAGLQFPVGRVHRLLRKGNYAERVGAGAPVYLAAVLEYLTAEILELAGNAARDNKKTRIIP |
| *Eriphia verrucosa* | 32 | RSSRAGLQFPVGRIHRLLRKGNYAERVGAGAPVYLAAVMEYLAAEVLELAGNAARDNKKTRIIP |
| *Eriocheir sinesis* | 9 | RSSRAGLQFPVGRIHRLLRKGNYAERVGAGAPVYLAAVMEYLAAEVLELAGNAARDNKKTRIIP |
| *Daphnia pulex* | 10 | RSSRAGLQFPVGRIHRMLRKGSYAERVGAGAPVYLAAVMEYLAAEVLELAGNAARDNKKTRIIP |
| *Macrobrachium nipponense* | 2 | RSSRAGLQFPVGRVHRLLRKGNYAERVGAGAPVYLAAVLEYLTAEILELAGNAARDNKKTRIIP |
| *Penaeus vannamei* | 42 | RSSRAGLQFPVGRIHRLLRKGNYAERVGAGAPVYLAAVMEYLAAEVLELAGNAARDNKKTRIIP |
| *Procambarus clarkii* | 11 | RSSRAGLQFPVGRIHRLLRKGNYAERVGAGAPVYLAAVMEYLAAEVLELAGNAARDNKKTRIIP |
| *Penaeus monodon* | 20 | RSSRAGLQFPVGRIHRLLRKGNYAERVGAGAPVYLAAVMEYLAAEVLELAGNAARDNKKTRIIP |
| *Scylla paramamosain* | 17 | RSSRAGLQFPVGRIHRLLRKGNYAERVGAGAPVYLAAVMEYLAAEVLELAGNAARDNKKTRIIP |
| *Portunus trituberculatus* | 14 | RSSRAGLQFPVGRIHRLLRKGNYAERVGAGAPVYLAAVMEYLAAEVLELAGNAARDNKKTRIIP |
| *Capitulum mitella* | 4 | RSSRAGLQFPVGRIHRLLRKGNYAERVGAGAPVYLAAVMEYLAAEVLELAGNAARDNKKTRIIP |
| *Palaemon carinicauda* | 41 | RSSRAGLQFPVGRIHRLLRKGNYAERVGAGAPVYLAAVMEYLAAEVLELAGNAARDNKKTRIIP |
| *Tigriopus californicus* | 11 | RSNRAGLQFPVGRIHRLLRKGNYAERVGAGAPVYLAAVMEYLAAEVLELAGNAARDNKKTRIIP |
| *Lepeophtheirus salmonis* | 6 | RSSRAGLQFPVGRIHRLLRKGNYAERVGAGAPVYLAAVMEYLAAEVLELAGNAARDNKKTRIIP |
| *Daphnia pulicaria* | 4 | RSSRAGLQFPVGRIHRMLRKGSYAERVGAGAPVYLAAVMEYLAAEVLELAGNAARDNKKTRIIP |
| *Coenobita brevimanus* | 117 | RSSRAGLQFPVGRIHRLLRKGNYAERVGAGAPVYLAAVMEYLAAEVLELAGNAARDNKKTRIIP |
| *Ceratothoa steindachneri* | 8 | RSSRAGLQFPVGRIHRLLRKGNYAERVGAGAPVYLAAVMEYLAAEVLELAGNAARDNKKTRIIP |
| *Morinoia aosen* | 23 | RSSRAGLQFPVGRIHRLLRKGNYAERVGAGAPVYLAAVMEYLAAEVLELAGNAARDNKKTRIIP |
| *Tethysbaena scabra* | 17 | RSSRAGLQFPVGRIHRLLRKGNYAERVGAGAPVYLAAVMEYLAAEVLELAGNAARDNKKTRIIP |
| *Pandarus bicolor* | 8 | RSSRAGLQFPVGRIHRLLRKGNYAERVGAGAPVYLAAVMEYLAAEVLELAGNAARDNKKTRIIP |
| *Jaera ischiosetosa* | 4 | RSNRAGLQFPVGRIHRLLRKGNYAERVGAGAPVYLAAVMEYLAAEVLELAGNAARDNKKTRIIP |
| *Anilocra frontalis* | 6 | RSSRAGLQFPVGRIHRLLRKGNYAERVGAGAPVYLAAVMEYLAAEVLELAGNAARDNKKTRIIP |
| *Jaera praehirsuta* | 5 | RSNRAGLQFPVGRIHRLLRKGNYAERVGAGAPVYLAAVMEYLAAEVLELAGNAARDNKKTRIIP |
| *Paralithodes platypus* | 62 | RSSRAGLQFPVGRIHRLLRKGNYAERVGAGAPVYLAAVMEYLAAEVLELAGNAARDNKKTRIIP |
| *Artemia sinica* | 13 | RSSRAGLQFPVGRIHRLLRKGNYAERVGAGAPVYLAAVMEYLAAEVLELAGNAARDNKKTRIIP |
| *Sacculina carcini* | 12 | RSSRAGLQFPVGRIHRLLRKGNYAERVGAGAPVYLAAVMEYLAAEVLELAGNAARDNKKTRIIP |
| *Daphnia carinata* | 6 | RSSRAGLQFPVGRIHRMLRKGSYAERVGAGAPVYLAAVMEYLAAEILELAGNAARDNKKTRIIP |
| *Cancer borealis* | 36 | RSSRAGLQFPVGRIHRLLRKGNYAERVGAGAPVYLAAVMEYLAAEVLELAGNAARDNKKTRIIP |
| *Daphnia galeata* | 1 | RSSRAGLQFPVGRIHRMLRKGSYAERVGAGAPVYLAAVMEYLAAEVLELAGNAARDNKKTRIIP |
| *Asellus aquaticus* | 5 | RSSRAGLQFPVGRIHRLLRKGNYAERVGAGAPVYLAAVMEYLAAEVLELAGNAARDNKKTRIIP |
| *Panulirus homarus* | 52 | RSSRAGLQFPVGRIHRLLRKGNYAERVGAGAPVYLAAVMEYLAAEVLELAGNAARDNKKTRIIP |
| *Panulirus ornatus* | 21 | RSSRAGLQFPVGRIHRLLRKGNYAERVGAGAPVYLAAVMEYLAAEVLELAGNAARDNKKTRIIP |
| *Acartia tonsa* | 11 | RSSRAGLQFPVGRIHRLLRKGNYAERVGAGAPVYMAAVMEYLAAEVLELAGNAARDNKKTRIIP |
| *Penaeus chinensis* | 40 | RSSRAGLQFPVGRIHRLLRKGNYAERVGAGAPVYLAAVMEYLAAEVLELAGNAARDNKKTRIIP |
